# Stepwise Oxidations Play Key Roles in the Structural and Functional Regulations of DJ-1

**DOI:** 10.1101/2021.01.25.428026

**Authors:** In-Kang Song, Mi-Sun Kim, James E. Ferrell, Dong-Hae Shin, Kong-Joo Lee

**Affiliations:** College of Pharmacy and Graduate School of Pharmaceutical Sciences, Ewha Womans University, Seoul, Korea 03760; Department of Chemical and Systems Biology, Stanford University School of Medicine, Stanford, Ca 94305-5174, USA

**Keywords:** DJ-1, redox-sensitive protein, cysteine oxidative modifications, intra-disulfide bond, hydrogen-deuterium exchange mass spectrometry (HDX-MS), stepwise oxidation

## Abstract

DJ-1 is known to play neuroprotective roles by eliminating reactive oxygen species (ROS) as an antioxidant protein. However, the molecular mechanism of DJ-1 function has not been well elucidated. This study explored the structural and functional changes of DJ-1 in response to oxidative stress. We found that Cys46 is also reactive cysteine residue in DJ-1, which was identified employing an NPSB-B chemical probe that selectively reacts with redox sensitive cysteine sulfhydryl. Peroxidatic Cys46 readily formed an intra-disulfide bond with resolving Cys53, which was identified with nanoUPLC-ESI-q-TOF tandem mass spectrometry (MS/MS) employing DBond algorithm under the non-reducing condition. We also found that Cys46-Cys53 disulfide crosslinking affects the oxidative state of the third Cys106, which shows the crosstalk among three cysteine residues of DJ-1. Furthermore, we demonstrated that DJ-1 C46A mutant, not forming Cys46-Cys53 intra-disulfide bond, lost structural stability of DJ-1 employing hydrogen/deuterium exchange-mass spectrometry (HDX-MS) analysis. All three Cys mutants lost antioxidant activities in SN4741 cell, a dopaminergic neuronal cell, unlike wild type DJ-1. These findings suggest that DJ-1 regulates its structure and activities by concerted oxidative modifications of three cysteine residues. These studies broaden the understanding of regulatory mechanisms of DJ-1 that operate under oxidative conditions.

## Introduction

Pathological feature of Parkinson’s disease (PD) is the selective degeneration of dopaminergic neurons, controlling key neurotransmitters related to movement, cognition, learning, and mood in the *Substantia Nigra pars compacta* (SNpc) of the human brain [1]. Mutations in several genes including DJ-1, α-synuclein, Parkin, Pink1, LRRK2 and UCH-L1 cause autosomal recessive PD [2] and increased ROS levels were observed in SNpc of sporadic PD patients [3]. Of these genes, several mutations in DJ-1 encoding the *PARK7* gene, are associated with early onset familial PD [4]. Since then, huge amounts of effort have been put to understand the contribution of DJ-1 to PD pathology. DJ-1 consisted of 189 amino acids, is evolutionary well conserved and ubiquitous small protein abundantly present in brain, especially in the *Substantia Nigra* [5]. Overexpression of DJ-1 results in neuronal cell protection against oxidative stress, while DJ-1 deficiency precedes oxidative stress induced cell death in cultured neuroblastoma cells and drosophila animal models [6–9]. In addition to antioxidant function of DJ-1, several different functions including transcriptional regulation and glyoxalase/deglycase activities have been assigned to DJ-1, which should be further studied for understanding the cause of these functions to PD [10].

L166P mutant of DJ-1, known mutant showing parkinsonian phenotype [11], is poorly folded [12] and lost its functions on eliminating free radicals and protecting cells [13], and oxidized DJ-1 is found in the brains of idiopathic PD individuals [14, 15]. So, structural changes of DJ-1 in response to mutations and oxidative stress have been intensively studied. DJ-1 has a flavodoxin-like Rossmann-fold containing a parallel β–sheet, and exists as a homodimer in crystal [16]. Its N-terminal sequence is necessary for the localization of DJ-1 to mitochondria [17]. The G-helix and kink portions of the C-terminal region are critical for the stability and redox-signaling of DJ-1 [18]. Analyses of the crystal structures of DJ-1 elucidate Cys106 residues at the sharp turn between a β-strand and an α-helix called the “nucleophile elbow”. Cys106 has energetically distorted angle, which raises reactivity of Cys106 by decreasing pKa [12] and by interacting with the protonated Glu18 residue *via* a hydrogen bond [19]. Oxidized DJ-1 at Cys106 to sulfenic and sulfinic acid and PD-associated known mutants (E64D, M26I, A104T, D149A) except L166P mutant demonstrate the modest structural changes compare to WT DJ-1 except subtle changes in weak chemical binding at residues [20–22]. In order to determine the dynamic structural changes of DJ-1 in response to oxidative stress, a hydrogen/deuterium exchange-mass spectrometry (HDX-MS) method was employed to fill a lack of dynamic changes absent in X-ray crystal structures [23]. In previous works with HDX-MS, we identified similar oxidative regulation in Nm23-H1 (NDPK-A) [24], peroxiredoxin 2 [25] and human secretagogin (hSCGN) by Ca^2+^ binding [26]. Hexameric Nm23-H1/NDPK-A, a tumor metastasis suppressor, is readily lost enzymatic activity by oxidizing Cys109 to sulfonic acid, even it is not active site, by oxidative stress [27], although Cys4 is the most redox sensitive residue. Single reactive cysteine can be easily oxidized and modified to be sulfenic acid (−SOH), sulfinic acid (−SO_2_H) and sulfonic acid (−SO_3_H). In addition, it contributes to form a disulfide bond together with adjacent resolving cysteine which is not reactive [28]. It has been found that reactive Cys4 forms intra-disulfide bond with Cys145, the resolving cysteine. It turns out that Cys4-Cys145 intra-disulfide bonds triggers a large conformational change that destabilizes the hexameric state which makes Cys109 oxidized to sulfonic acid. C4A and C145A mutants cannot oxidize Cys109 and are always active as well as C109A [24]. This indicates stepwise oxidation is one way of oxidative regulation of proteins. In addition to stepwise oxidation, oxidized proteins can be readily degraded in proteasome by oxidation induced structural change makes the protein be ubiquitinated in redox sensitive proteins including peroxiredoxin and DJ-1 etc [25]. Secretagogin, a hexa EF-hand calcium binding protein functioning in insulin secretion in pancreatic beta-cells, induces the structural change by calcium binding, which triggers the formation of functional dimer via inter-disulfide crosslinking [26]. These examples suggest the stepwise regulation combining oxidation with oxidation, ubiquitination and phosphorylation [29].

DJ-1 is a redox-sensitive protein easily oxidized and degraded by oxidative stress [25]. Because antioxidant proteins have reactive Cys residues with redox capacity, we focused the property of cysteine residues in DJ-1. There are three cysteines in DJ-1; Cys46, Cys53 and Cys106. Orthologous analysis demonstrated that Cys106 [30] and Cys46 are well conserved in every species from *C. elegans* to *H. sapiens*. However, Cys53 is conserved in mammalian only (Fig. 1A, Supplementary Table 1). In general, evolutionary well-conserved residues have a tendency to be transmitted to a lineage of life and can play important roles in effective protein folding, stability and function [31]. Of these, Cys106 is known as the most readily modified residue by oxidation to sulfinic acid [30, 32–34], which shifts DJ-1 to the mitochondria, thereby protecting neurons from oxidative stress. To date, most research on DJ-1 has been focused on Cys106, but recently the importance of Cys46 and Cys53 has also been raised. Cys46 and Cys53, but not Cys106, are susceptible to S-nitrosylated and Cys46 is involved in the dimerization of DJ-1[35]. Cys53 and Cys106 are known to react with dopamine to form adducts [36], and Cys53 are known to be needed for dimer formation because they are near the dimer interface [37]. However, it is not still possible to explain how DJ-1 can sense oxidative stress, how DJ-1 protects cells from oxidative stress, and how DJ-1 contribute PD pathogenesis. Here, we focus how DJ-1 is modified in response to oxidative stress and how Cys mutants induce the structural changes and regulate the cellular functions.

**Fig. 1.**
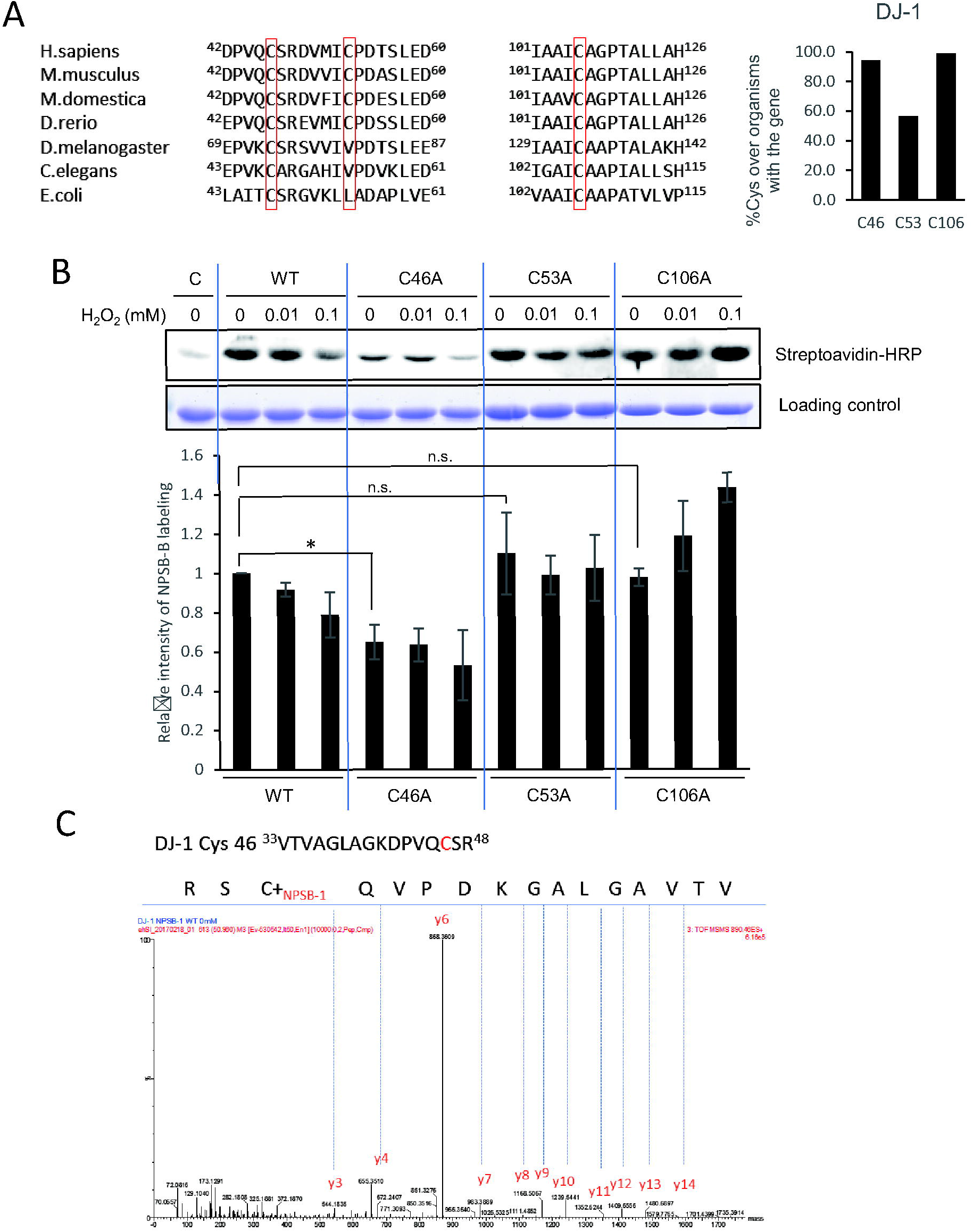
Cys46 is the most reactive residue in DJ-1. (A) Alignment of amino acid sequences in DJ-1. Cys46 residue and Cys106 residue are well-conserved. (B) Recombinant WT and Cys mutant DJ-1 proteins were incubated with indicated concentrations of H_2_O_2_ at 37°C for 1 h followed by treatment with 1 mM NPSB-B at RT for 2 h. Proteins were separated on reducing SDS-PAGE and detected by streptavidin-HRP. Coomassie staining gel showing amount of gel loaded proteins. NPSB-B labeling were quantified and normalized with loading control. Data are presented as mean ± S.D. of triplicated experiments (Two tailed t-test; * *P* < 0.05). (C) Tandem mass spectrum of NPSB-1 labeled at Cys46 residue in WT DJ-1. WT protein was tested in same way as NPSB-B, separated by SDS-PAGE, and stained with Coomassie blue, then labeled bands were cut out of gel and analyzed by MS, and PTM analysis was performed.

In the present study, using a novel specific biotin-labeling chemical probe, which reacts with redox-sensitive Cys-SH residues [38] and HDX-MS [23], we identified that the reactive Cys residue in DJ-1 is Cys46, and found that C46A mutant display profound structural defects like L166P DJ-1 mutant. We focused on oxidative modifications and structural changes of three DJ-1 Cys mutants and suggest that Cys46 residue is critical to maintain stable structure and Cys46-Cys53 intra-disulfide bond regulates the oxidative status of Cys106 in concerted way. We further demonstrated that all three DJ-1 mutants (C46A, C53A, C106A) abolished the protective functions against oxidative stress in neuronal cell. These results reveal that all three cysteine residues of DJ-1 are indispensable for maintaining the structure, regulation process and function as an antioxidant protein.

## Materials and Methods

### Cell culture and transfection

Human cervical carcinoma HeLa cells from ATCC (VA, USA) were cultured in Eagle’s minimum essential medium (EMEM) supplemented with 10% fetal bovine serum (FBS), 100 μg/mL streptomycin and 100 units/mL penicillin G at 37°C in an atmosphere of 5% CO_2_ −95% air. SN4741 dopaminergic neuronal cell was cultured at 33°C with 5% CO_2_ in RF medium containing DMEM supplemented with 10% FBS, 1% glucose, 100 μg/mL streptomycin and 100 units/mL penicillin G, and L-glutamine [47]. For transient overexpression of specific proteins in HeLa cells and SN4741 DJ-1 KO cells, cells were transfected using LT-1 and analyzed 48 h later.

### Plasmids

Human DJ-1 were cloned into pFlag-CMV2 and pGEX-4T-1 vectors. Cys to Ala mutants of Flag-tagged DJ-1 were using QuikChange Ⅱ Site-Directed Mutagenesis kit (Agilent Technologies, USA) according to the manufacturer’s protocol. The primers used for mutagenesis are as follows: C46A_s, 5’-GAA AAG ACC CAG TAC AGG CTA GCC GTG ATG TGG TC-3’; C46A_as, 5’-GAC CAC ATC ACG GCT AGC CTG TAC TGG GTC TTT TC-3’; C53A_s, 5’-CCG TGA TGT GGT CAT TGC TCC TGA TGC CAG CCT TG-3’; C53A_as, 5’-CAA GGC TGG CAT CAG GAG CAA TGA CCA CAT CAC GG-3’; C106A_s, 5’-CCT GAT AGC CGC CAT CGC TGC AGG TCC TAC TGC TC-3’; C106A_as, 5’-GAG CAG TAG GAC CTG CAG CGA TGG CGG CTA TCA GG-3’. All plasmids were confirmed by DNA sequencing.

### Purification of Recombinant proteins

A pGEX-4T-1 plasmid carrying human DJ-1 gene was expressed in BL21 (DE3) *E.coli* cells. Bacteria were cultured in LB medium at 37°C, and recombinant fusion protein production was induced with 0.25 mM were isopropyl-β-D-thiogalactopyranoside (IPTG). After 4 h of additional incubation at 37°C, the cells were harvested and lysed by vortexing with lysozyme containing lysis buffer (lysozyme, protease inhibitor, triton X-100 in PBS (140 mM NaCl, 2.7 mM KCl, 10 mM Na_2_HPO_4_, pH 7.4)) on ice and sonicated. Soluble protein fraction was recovered by centrifugation at 13,000 g for 30 min at 4°C. GST-fused recombinant proteins in supernatant were purified by chromatography on a glutathione-agarose column followed by washing, and equilibration by elution buffer (140 mM NaCl, 2.7 mM KCl, 10 mM Na_2_HPO_4_, pH 7.4). To cleave DJ-1 from GST-DJ-1, the beads were incubated overnight at RT with thrombin in DJ-1 elution buffer. After 16 h, purified DJ-1 was eluted and protein concentration was quantified with the BCA protein assay.

### NPSB-B labeling

Methods for the labeling of reactive cysteine residues by NPSB-B were previously reported [38]. Briefly, 2 μg of recombinant DJ-1 was pre-incubated with various concentration of H_2_O_2_ at 37°C for 1 h followed by incubation with NPSB-B (final concentration 1 mM) at RT for 2 h. Proteins were separated by 12% SDS-PAGE and transferred to PVDF membranes. Labeled proteins were detected by streptavidin-HRP. Amount of total loaded protein was detected by Coomassie staining.

### Identification of Cys modifications employing nanoUPLC-ESI-q-TOF tandem MS

Recombinant WT and mutant proteins (2 μg) and over-expressed Hela cell with DJ-1 WT and mutants were incubated with various concentrations of H_2_O_2_ at 37°C for 1 h followed by separation by SDS-PAGE under non-reducing and reducing condition. In order to prevent oxidation during sample preparation, alkylating agent N-ethylmaleimide (NEM) was treated in several indicated experiments. All experiments were triplicated. Gel bands of differentially expressed proteins were destained and digested with trypsin, and the resulting peptides were extracted as previously described [58]. Peptides were separated using trap column cartridge, injected into C18 reversed-phase analytical column with integrated electrospray ionization PicoTip™ using nanoAcquity™ UPLC/ESI/q-TOF MS/MS (SYNAPT™ G2Si; Waters Co.). Peptide solutions were injected into column and eluted by linear gradient of 5–40% buffer B (ACN/formic acid; 100:0.1, v/v) with buffer A (Water/formic acid; 100:0.1, v/v) over 60 min. Mass spectrometer was programmed to record scan cycles composed of one MS scan followed by MS/MS scans of the 10 most abundant ions in each MS scan.

### Search parameters and acceptance criteria for MS data

Pkl files were generated using ProteinLynx global server 2.3 (Waters Co.,). Peaklists were searched against DJ-1 FASTA file (Uniprot accession no. Q99497) using Mascot (version 2.2.06, Boston, USA), MOD^i^ [59] and Modplus (https://prix.hanyang.ac.kr). For Cys oxidative modification search of samples, oxidation, dioxidation and trioxidation of Cys, Cys-SO_2_-SH, conversion of Cys to Ser, and acrylamide adduct propionamide of Cys were used as variable modifications. Mass tolerance for precursor ions was ± 0.5 Da and mass tolerance for fragment ions was ± 0.5 Da because the error value of MS was about 0.01. Significant matches were sorted by the threshold indicated by Mascot probability analysis, and matches with probability-based Mowse score P < 0.05 (and expectation value of 0.05) were considered significant. Ions score was calculated as −10 x Log (P) (P: the probability that the observed match is a random event) and automatically provided as Mascot result. Regarding the protein identification result, the best match with the highest score amongst identified results was used as the identification result for the spot. PTM results were verified not just by score threshold, but by checking each spectrum carefully. MS peaks in Supplementary Fig. 3 were generated by MOD^i^ spectrum viewer [59].

### Disulfide analysis of DJ-1 using nanoUPLC-ESI-q-TOF tandem MS and DBond algorithm

Peptides resulting from trypsin-digested protein (for C46A, Glu-C and trypsin-) in non-reducing gel were resuspended in 10% acetonitrile containing 0.1% formic acid and analyzed using nanoAcquity™ UPLC™/ESI/MS (SYNAPT™ G2Si, Waters Co. UK) as described previously [58]. Tandem MS (MS/MS) spectra were matched against DJ-1 fasta (Uniprot accession no. Q99497) using DBond algorithm (https://prix.hanyang.ac.kr) [40]. For recombinant protein sample, mass tolerance for precursor ions was ± 0.2 Da and mass tolerance for fragment ions was ± 0.2 Da and for Hela cell sample, mass tolerance for precursor ions was ± 0.5 Da and mass tolerance for fragment ions was ± 0.5 Da due to low intensity). DBond score was calculated by byIonScore, bondScore and sulfurScore and automatically provided as DBond result. For recombinant protein sample, the sample intensity was high, only scores above 50 were taken. MS peaks in Figs. 2B, 6C, Supplementary Fig. 3 and 6 were generated by MOD^i^ spectrum viewer [59].

**Fig. 2.**
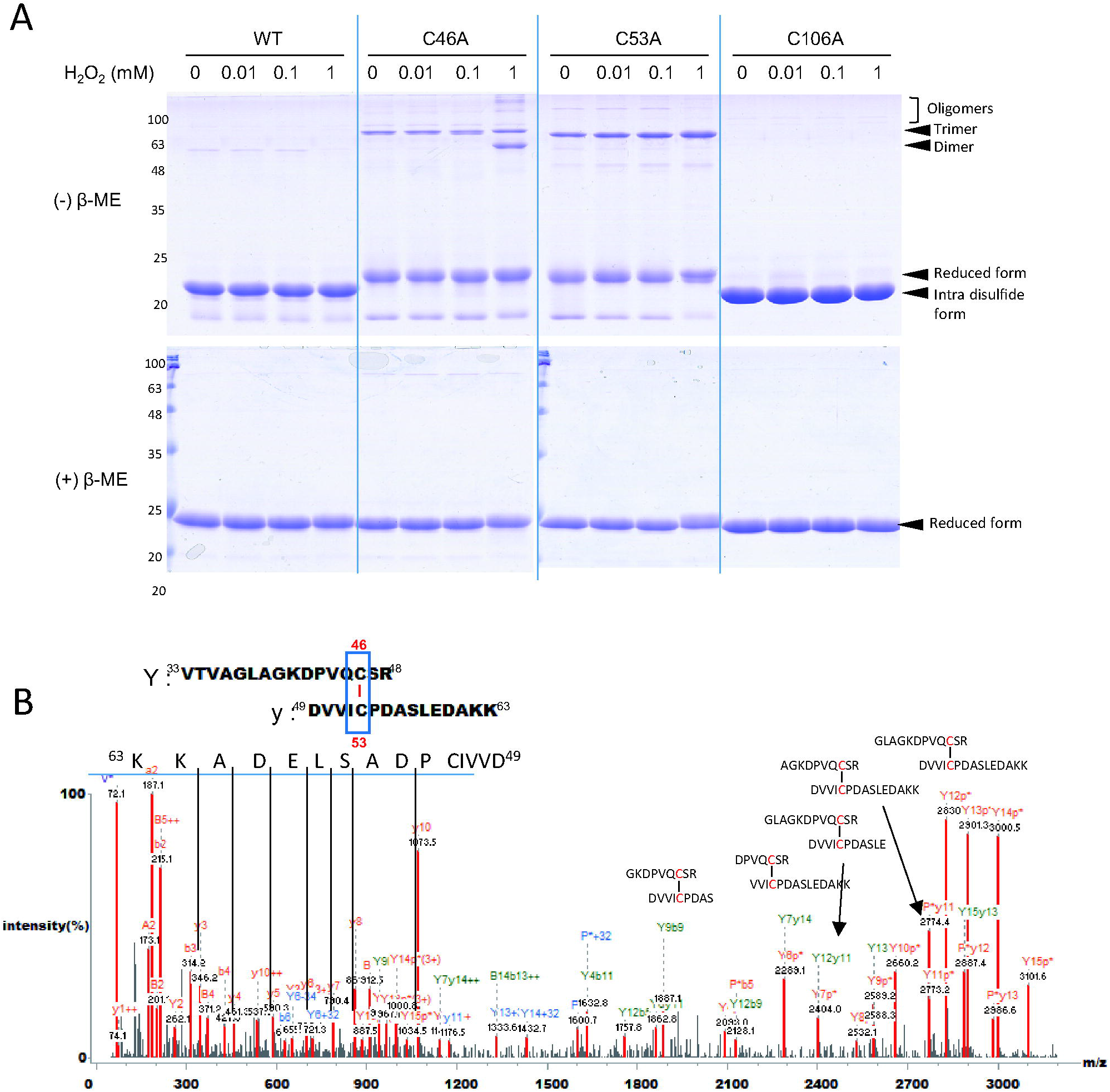
DJ-1 has a Cys46-Cys53 disulfide bond in recombinant proteins. (A) Purified WT and Cys mutants DJ-1 proteins were incubated with 0, 0.01, 0.1 or 1 mM H_2_O_2_ at 37°C for 1 h. Proteins were separated by SDS-PAGE under non-reducing (−β-ME) and reducing (+β-ME) conditions, and detected by Coomassie blue-staining. (B) Tandem mass spectrum of Cys46-Cys53 disulfide bond peptide of WT DJ-1 band without H_2_O_2_ treatment in Fig. 2A. Coomassie-stained gel bands were cut out of gel and analyzed by MS and Dbond algorithm.

### HDX-Hydrogen/deuterium exchange mass spectrometry (HDX-MS)

Methods for the HDX-MS were previously reported [26]. DJ-1 protein (1 μg) was diluted 9-fold with labeling buffer (10 mM HEPES in D_2_O, pH 7.4) and incubated at 25°C for 10, 60, 300, 1800, or 10800 s. Deuterium labeling reaction was quenched by 2.5 mM tris (2-carboxyethyl) phosphine (TCEP), formic acid, pH 2.3. For protein digestion, 1 μg of porcine pepsin was added to each quenched protein sample and incubated on ice for 3 min before injection. Peptic peptides were desalted on C18 trap column cartridge (Waters, UK) and gradient eluted from 8% CH_3_CN to 40% CH_3_CN, 0.1% formic acid on 100 μm i.d. × 100 mm analytical column, 1.7 μm particle size, BEH130 C18, (Waters, UK) for 7 min. The trap, analytical column and all tubing were immersed in ice bath to minimize deuterium back-exchange. Gradient chromatography was performed at a flow rate 0.5 μL/min and was sprayed on line to nanoAcquity™/ESI/MS (SYNAPT™ HDMS™) (Waters, UK). The extent of deuterium incorporation was monitored by increase in mass of isotope distribution for each identified peptide, and calculated using Microsoft Excel. Each experiment was performed in triplicate.

### Circular dichroism (CD)

Circular dichroism (CD) and thermal denaturation were recorded in buffer containing 137 mM NaCl, 2.7 mM KCl and 10 mM Na_2_HPO_4_ at pH 7.4 using a JASCO J-1500 spectropolarimeter (Jasco, Tokyo, Japan). The concentration of protein solution was 0.25 mg/mL for CD experiments. Monitoring of protein thermal denaturation was performed in temperature range 20-90°C, increasing the temperature at the rate of 1°C/min. After 1 min of equilibration at each temperature, ellipticity was measured at 222 nm.

### Protein denaturation induced by GdnHCl

Protein unfolding induced by guanidinium hydrochloride (GdnHCl) was performed by incubating WT and mutants proteins in PBS containing various concentrations of GdnHCl (pH 7.4) ranging from 0 to 2.0 M 16 h at 25°C. Final protein concentration was 0.2 mg/mL. Unfolded samples were incubated with 8-anilinonaphtalene-1-sulfonic acid (ANS) for fluorescence measurements to monitor structural changes induced by unfolding.

### Co-immunoprecipitation

HeLa cells were seeded at a density of 7.5 × 10^5^ cells/100-mm plate and grown for 24 h. Control plasmid pFlag CMV vector, and plasmids carrying wild-type (WT) and Cys mutants (C46A, C53A and C106A) of DJ-1 were delivered into cells respectively using LT-1 transfection reagent. At 24 h post-transfection, cells were washed twice with 2 mL Hanks’ Balanced Salt Solution (HBSS) to remove serum. Cells were treated with 0 or 0.5 mM of H_2_O_2_ in HBSS for 1 h (37°C, 5% CO_2_), then lysed in IP lysis buffer (50 mM Tris-Cl, pH 7.4, 150 mM NaCl, 1 mM EDTA, 0.5% NP40, pH 7.4) supplemented with protease inhibitor mixture (Sigma) and phosphatase inhibitors (2.5 mM Na_3_VO_4_ and 2.5 mM NaF). Cells were passed 31-gauge syringe 10 times and centrifuged at 12,000 rpm for 10 min. Supernatant was immunoprecipitated with anti-Flag antibody for 2 h and then with protein G Sepharose beads for 1 h at 4°C. Beads were washed three times with 1 mL of lysis buffer and additionally twice with 1 mL of lysis buffer without detergent. Immune complex was solubilized in SDS gel sample buffer, separated on SDS-PAGE, and analyzed by silver staining and Western blot analysis.

### DCF-DA staining

Cellular ROS levels were measured using fluorescent dye, 2’, 7’ - dichlorodihydrofluorescein diacetate (CM-H_2_DCFDA) (Molecular Probes: OR, USA). SN4741 DJ-1 KO cells were incubated with 0 mM or 0.5 mM H_2_O_2_ in HBSS at 33°C for 20 min. For flow cytometry, cells were washed with cold PBS, and treated with 2 μM CM-H_2_DCFDA in HBSS at 33°C for 15 min in dark. Cells were collected by trypsinizing briefly and incubated in ice as pellet. The pellets was straightly suspended into buffer, and cellular ROS was immediately measured using FACSCalibur flow cytometer (BD Biosciences: NJ, USA). 10,000 cells were analyzed for each sample. To quantify log-amplified fluorescence which is emitted by cells, geometric mean (Geo Mean) of fluorescence intensity as well as median value for fluorescent peak was calculated by statistical analysis of BD CellQuest software.

### Antibodies

Antibodies used in this study were: Flag (Sigma), β–actin (Santa Cruz), pERK(T202/Y204), ERK (Abcam) and DJ-1 (Ab frontier).

### Cell proliferation assay

DJ-1 KO SN4741 cells were transiently transfected with Flag-DJ-1 WT and Cys mutants were incubated at 0 or 0.5 mM H_2_O_2_ 2 h at 33°C. Cells were trypsinized and seeded at a density 2,500 cells in 96-well RTCA E-plate. For real-time cell analyzer analysis, cell growth was real time monitored under xCELLigence RTCA SP system (Acea Biosciences, Inc.).

## Results

### Cys46 of DJ-1 is the reactive cysteine residue

In order to investigate the properties of Cys46 and Cys53 together with already well-known Cys106, the reactivity of these cysteines was examined using recombinant WT and Cys mutant proteins. They were incubated with various concentrations of H_2_O_2_ in 1 h and then labeled with NPSB-B, a novel specific biotin-labeling chemical probe, which reacts with redox-sensitive Cys-SH residues [38]. As shown in Fig. 1B, WT DJ-1 has a highly reactive cysteine residue, which is readily labeled with NPSB-B. The labeling decreased after oxidation in a H_2_O_2_-dose-dependent manner. The result indicates the presence of redox sensitive cysteine in DJ-1. The labeling result with C53A and C106A mutants also showed that the level of labeling with NPSB-B on both mutants was comparable with that of WT. In contrast, C46A mutant could not be well labeled with NPSB-B. The result indicates that Cys46 is the most redox sensitive Cys residue like Cys106. However, there is an increased labeling level detected on Cys106 mutant at 0.1 mM H_2_O_2._ This indicates that the reactivity of Cys46 increases as the concentration of H_2_O_2_ increases in the absence of C106, which requires further study. The result was corroborated by detecting labeled peptides at Cys46 with NPSB-1 which is a non-biotin probe with combining nanoUPLC-ESI-q-TOF mass spectrometry with Mascot algorithm. Tandem mass spectrum of the identified NPSB-1 labeling in Cys46 residue of WT DJ-1 (^33^VTVAGLAGKDPVQCSR^48^ (m/z=890.4360 Da, z = 2, Δ= +179.02 Da)) is shown in Fig. 1C. These results suggest that Cys46 residue is the most reactive cysteine in DJ-1.

### Cys46 and Cys53 form an intra-disulfide bond *in vitro*

Since highly reactive Cys46 does not undergo various oxidation [39], it is possible to form a disulfide bond. To investigate whether Cys46 could form a disulfide bond, we examined the electrophoretic mobility of WT and mutant DJ-1s on SDS-PAGE with or without 2-mercaptoethanol (β-ME) (Fig. 2A). WT and the C106A mutant were down-shifted under non-reducing conditions and returned to their original position under reducing conditions. The result indicated the presence of the same intra-disulfide bond in WT and C106A mutant considering the consistent pattern depending on H_2_O_2_ concentration. On the other hand, C46A and C53A mutants were mainly in the same position corresponding to a monomeric form in SDS-PAGE both under reducing and non-reducing conditions. In order to confirm that the down-shifted band formed an intra-disulfide bond, WT and C46A mutant were incubated with a reducing agent, 1,4-Dithiothreitol (DTT), by concentration, and only WT was reduced as the DTT concentration increased, confirming that intra-disulfide was released (Supplementary Fig. 1). These results suggested that C46A and C53A mutants do not form an intra-disulfide bond. It is worthwhile to mention that C46A mutant also forms a trimeric form and further aggregates with even higher molecular weight under 1 mM H_2_O_2_. C53A mutant was found to form a trimer of which the amount is proportional to H_2_O_2_ concentration.

In order to validate these results with direct recognition of intact disulfide bonds, we identified the disulfide bond in DJ-1 employing nanoUPLC-ESI-q-TOF mass spectrometry and DBond, a disulfide searching algorithm [40]. Tandem MS spectra have confirmed the presence of intra-disulfide bond between Cys46 and Cys53 in the main band of WT sample after tryptic digestion (^33^VTVAGLAGKDPVQCSR/DVVICPDASLEDAKK^63^ (m/z = 800.9005 Da, z = 4)) (Fig. 2B, Table 1). Intriguingly, C46A and C53A mutants lacking Cys46-Cys53 disulfide bond showed new disulfide bonds formed by Cys53-Cys106 (Supplementary Table 2) and Cys46-Cys106 (Supplementary Fig. 2, Table 1), respectively under oxidative stress, although scores are low in oxidized samples. The results show that DJ-1 is oxidized to form intra-disulfide bond primarily by Cys46 with Cys53, but three cysteine residues also seemed to influence each other’s oxidation states.

**Table 1.** List of identified intra-disulfide bonds in WT and mutant DJ-1 recombinant protein bands under the non-reducing condition in Fig. 2A. Each protein band was digested with trypsin and analyzed with MS combining with DBond algorithm. MS/MS spectra are presented in Fig. 2B and Supplementary Fig. 2.

| | H<sub>2</sub>O<sub>2</sub><br>(mM) | Disulfide<br>bond | Mass(m/z)<br>Experimental<br>(Charge state) | Mass<br>Theoretical | $\Delta m$ (Da) | Dbond<br>Score | Sequence |
| --- | --- | --- | --- | --- | --- | --- | --- |
| WT | 0 |  | 601.7895(4+) | 2403.141 | -0.0121 | 58.6 | DPVQC*SR-DVVIC*PDASLEDAKK |
|  | 0.01 | Cys46- | 800.9005(4+) | 3199.6216 | -0.0487 | 266.3 | VTVAGLAGKDPVQC*SR-DVVIC*PDASLEDAKK |
|  | 0.1 | Cys53 | 800.9019(4+) | 3199.6216 | -0.0431 | 390.4 | VTVAGLAGKDPVQC*SR-DVVIC*PDASLEDAKK |
|  | 1 |  | 800.902(4+) | 3199.6216 | -0.0427 | 313.7 | VTVAGLAGKDPVQC*SR-DVVIC*PDASLEDAKK |
| C46A | 0 |  |  |  |  |  | N.D. |
|  | 0.01 | Cys53- |  |  |  |  | N.D. |
|  | 0.1 | Cys106 |  |  |  |  | N.D. |
|  | 1 |  |  |  |  |  | N.D. |
| C53A | 0 |  |  |  |  |  | N.D. |
|  | 0.01 | Cys46- | 808.054(3+) | 2421.2144 | -0.0742 | 90.8 | DPVQC*SR-GLIAAIC*AGPTALLAHEIGFGSK |
|  | 0.1 | Cys106 |  |  |  |  | N.D. |
|  | 1 |  |  |  |  |  | N.D. |
| C106A | 0 |  | 800.9137(4+) | 3199.6216 | 0.0041 | 310.8 | VTVAGLAGKDPVQC*SR-DVVIC*PDASLEDAKK |
|  | 0.01 | Cys46- | 803.344(2+) | 1604.7035 | -0.0301 | 140.1 | VTVAGLAGKDPVQC*SR-DVVIC*PDASLEDAKK |
|  | 0.1 | Cys53 | 768.8725(4+) | 3071.5267 | -0.0658 | 628.5 | VTVAGLAGKDPVQC*SR-DVVIC*PDASLEDAK |
|  | 1 |  | 759.3374(3+) | 1604.7035 | -0.0406 | 193.8 | DPVQC*SR-DVVIC*PDASLEDAK |

\*N.D.: modified peptide were not detected

### Each cysteine influenced on the oxidation state of other cysteines with different modifications

The influence among three cysteines of DJ-1 has been investigated and analyzed by examining oxidative modifications of Cys residues in WT and three Cys mutants employing nanoUPLC-ESI-q-TOF tandem MS combining Mascot search algorithm. Sulfhydryl modifications such as sulfenic, sulfinic and sulfonic acids together with various oxidative conversion of cysteine to dehydroalanine (DHA), serine or Cys-SO_2_-SH resulting from breakdown of newly discovered thiosulfonate [41, 42] have been thoroughly examined in monomeric bands of H_2_O_2_ untreated samples appeared on reducing SDS-PAGE in Fig. 2A. The list of peptides containing each cysteine modification is shown in Fig. 3A. Cys106 was oxidized and modified into various oxidative forms (Fig. 3B); sulfinic acid (−SO_2_H, Δm = +32 Da), sulfonic acid (−SO_3_H, Δm = +48 Da) and thiosulfonic acid (−SO_2_-SH, Δm = +64 Da). In WT, Cys106 was detected as sulfinic acid, an active form found in many cysteine residues. However, Cys106s in C46A and C53A mutants were more oxidized to sulfinic acid and significantly further oxidized to sulfonic acid, an irreversibly oxidized form. These results indicate that Cys106s in C46A and C53A mutants are more oxidized than that in WT.

**Fig. 3.**
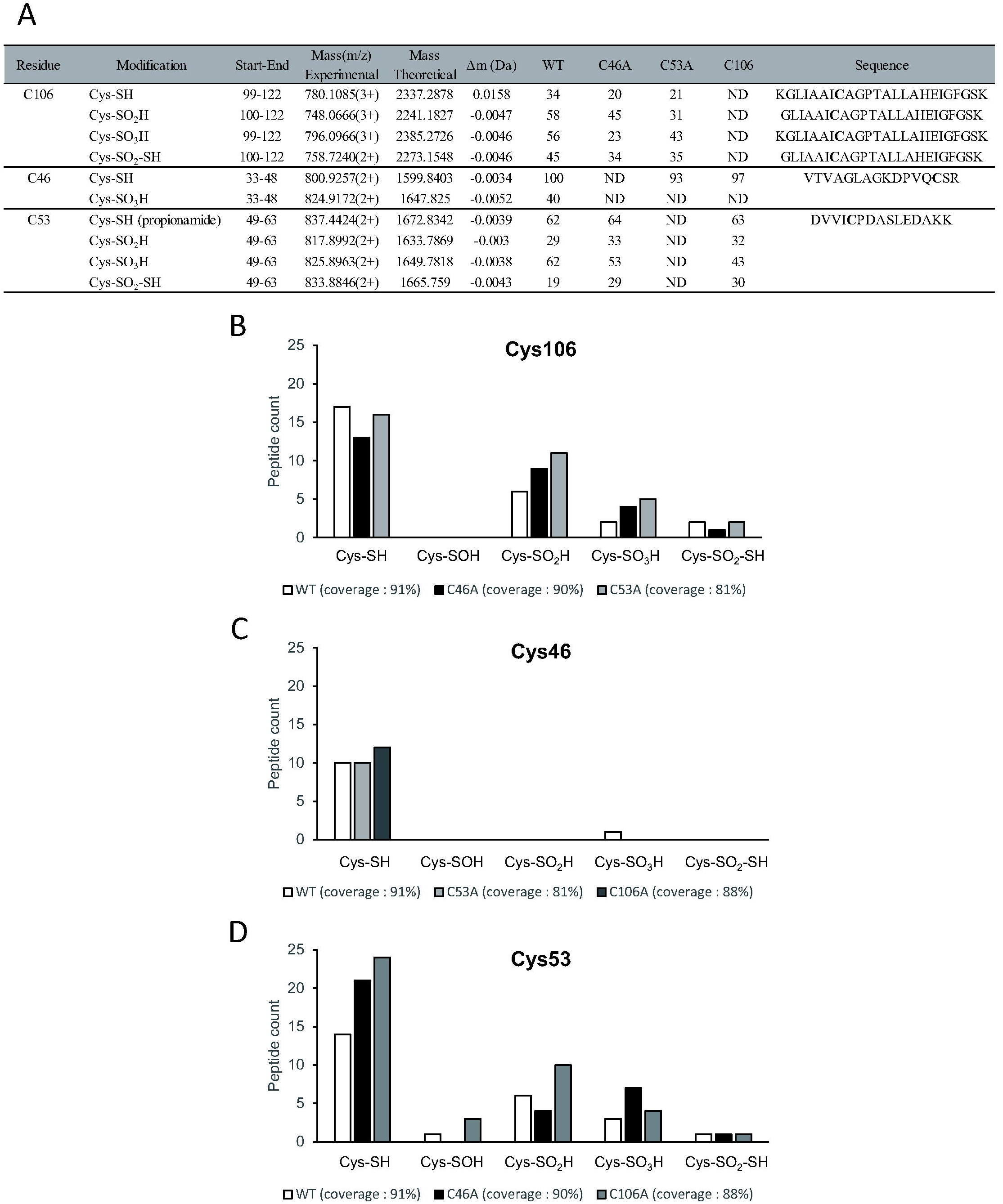
Each cysteine influences the oxidation state of other cysteines with different modifications. (A-D) Oxidative modifications of WT and Cys mutants DJ-1 protein bands in Fig 2A without H_2_O_2_ treatment under reducing condition were examined by MS/MS analysis employing SEMSA strategy and Mascot search algorithm. MS/MS spectra of modified peptides are presented in Supplementary Fig. 3. Numbers of oxidized peptides in (B) Cys106, (C) Cys46 and (D) Cys53 residue in WT and Cys mutant DJ-1s were presented with mass coverage. Various oxidative modifications such as sulfinic acid, sulfonic acid and conversion of Cys to thiosulfonic acid (Δm = +64 Da) were found.

Cys46 residues in WT, C53A and C106A are not oxidized except only one sulfonic acid in WT (Fig. 3C). Cys53s were slightly oxidized in WT, but more oxidized to sulfenic, sulfinic and sulfonic acid in C106A and further oxidized to sulfonic acid in C46A than those in WT (Fig. 3D). The results indicate that Cys106 is kept in sulfinic acid in WT, but is more oxidized to sufonic acid in mutant not forming disulfide bond. This suggests that one Cys mutation affects oxidative modifications of the other Cys residues.

### Intra-disulfide bond between Cys46-Cys53 in DJ-1 plays an important role in its structure

Consistent with the results from the above experiments, Cys46 and Cys53 appear at a distance of 11.54 Å in the X-ray crystal structure of human DJ-1 treated with DTT [43]. However, since both are on a flexible loop, it is possible they crosslink to form an intra-disulfide bond. (Fig. 4C). In the case of C46A mutant, a large conformational change should be happened to form Cys53-Cys106 intra-disulfide bond (Supplementary Table 2) due to a distant location of Cys53 and Cys106 with a distance of 23.08 Å. To investigate the influence of Cys46-Cys53 intra-disulfide bond on structural stability of DJ-1, we conducted HDX-MS analysis of WT and C46A mutant. We mapped DJ-1 sequence onto the structure deduced from X-ray crystallographic studies of *H. sapiens* DJ-1 (PDB ID code 4KRW) with chimera [44] and used this structure to display the results of the HDX-MS studies. H/D exchange rates of WT and C46A mutant were compared in each peptic digested peptide (Fig. 4A, Supplementary Table 3). Combined stitching H/D exchange ratios of peptic peptide showed the diagram of whole protein shown in Fig. 4B and overlay of differential HDX data between WT and C46A mutant onto the structures of human DJ-1 in Fig. 4C. The peptide MS coverage was 100%. This study revealed the differences between WT and C46A mutant with respect to deuterium incorporation rates in specific regions of protein. In most areas, C46A mutant exhibited higher deuterium exchange than WT. This indicates that C46A mutant is structurally dynamic and more exposed to solvent. In particular, the peptide comprising Cys106 residue (a.a 104-112) identified as involved in active site had higher deuterium exchange. As well as, deuterium exchange levels of peptides nearby Cys106 in tertiary structure also increased in C46A mutant. Cys106 is adjacent to Glu18 and His126. The former depresses pKa of Cys106 and modulates the reactivity of Cys106 [19], and the latter is member of catalytic diad with Cys106 in DJ-1 superfamily [34]. In HDX-MS analysis, peptides containing regions around Glu18 (a.a.17-26) and His126 (a.a 117-131) in C46A mutant significantly increased deuterium exchange levels. These results indicate that surrounding regions of Cys106 in C46A mutant are dramatically exposed to the protein surface, which is consistent with above proteomic results that Cys106 residue in C46A mutant is further oxidized to sulfonic acid.

**Fig. 4.**
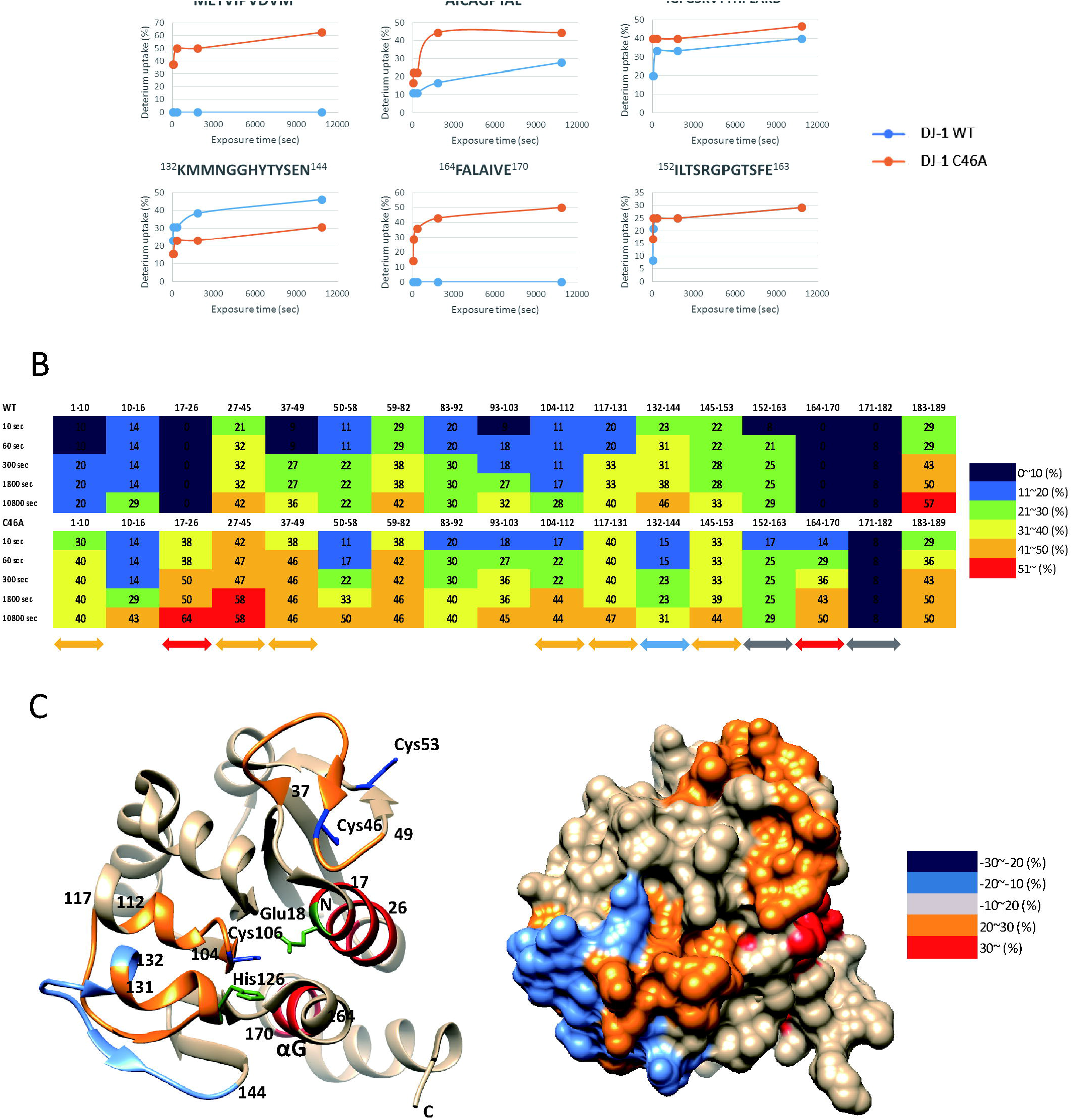
Identification of structural changes in WT and C46A mutant DJ-1 employing HDX-MS. (A-C) Recombinant WT and C46A mutant DJ-1 proteins were incubated with D_2_O exchange buffer at 25°C for various times upto 30 min and analyzed using nanoAcquity™/ESI/MS. (A) Time course of HDX incorporation for representative peptides that showed differences in HDX. (B) Deuterium exchange rate (%) of each peptide in WT and C46A mutant was presented depending on D_2_O incubation time. Significant differences between WT and C46A mutant were marked by colored arrows and no discernible changes by gray arrow. Corresponding deuterium exchange levels of each peptide in percent are given on the right. (C) Average deuterium exchange difference of C46A mutant compared to the WT in DJ-1 monomer structure (PBD ID code 4RKW).

Additionally, the region near Ala46 (a.a 37-49) in C46A mutant also had increased exposure, indicating that Cys46-Cys53 intra-disulfide bond were released and became more flexible. The peptide of αG (a.a 164-170) in helix-kink-helix motif which is essential for protein stability [18] also showed the significantly higher exchange level in C46A mutant. Meanwhile, the peptide (a.a 132-144) in the opposite side of Cys106 was more shielded in the mutant. And, the peptide (a.a 152-163) in helix of C-terminus showed similar exchange level in both WT and mutant. On the other hand, as a result of HDX-MS analysis of oxidized WT, there was no significant difference from native WT except that the peptide including Cys46 was shielded and the peptide including Cys106 was increased (Supplementary Fig. 4). In summary, the results of these dynamic structural analyzes indicate that C46A mutant, not forming the intra-disulfide bond, undergoes significant conformational changes.

### C46A mutant is more unstable than WT *in vitro*

To investigate protein stability of C46A mutant, not able to form Cys46-Cys53 intra-disulfide bond, we examined protein stability of WT and C46A mutant employing biophysical assessments of denaturation induced by heat or chemical denaturants. We measured CD ellipticity of WT and C46A mutant (0.25 mg/mL) at 222 nm by increasing temperature (20-90°C) at a rate of 1°C/min. As shown in Fig. 5A, mid-point of thermal transitions, *T*_m_, were 61.3°C for WT and 51.5°C for C46A mutant. A dramatic destabilization effect was observed in C46A mutant which led to 9.8°C decrease in *T*_m_. Thus, WT has more stable structure than C46A mutant. In addition, protein folding was examined using the chemical denaturant, GdnHCl. Protein unfolding was investigated by measuring ANS fluorescence after treatment of various concentrations of GdnHCl using the properties of accumulating ANS bound to unfolded hydrophobic region of protein [45]. ANS fluorescence intensity of WT was increased at 0.6 M GdnHCl concentration, while that of C46A mutant was rapidly increased at 0.3 M (Fig. 5B). This indicates that C46A mutant is more readily denaturized than WT. These results imply that the conformational change in C46A mutant may expose more hydrophobic areas on its molecular surface. Additionally, C46S, a conserved mutant, was confirmed to be unstable like C46A (Supplementary Fig. 5A), and C53A mutant, which does not form an intra-disulfide bond, was not as unstable as C46A, but more unstable than WT (Supplementary Fig. 5B). These results indicate that structure of C46A mutant is distorted due to the lack of Cys46-Cys53 intra-disulfide bond.

**Fig. 5.**
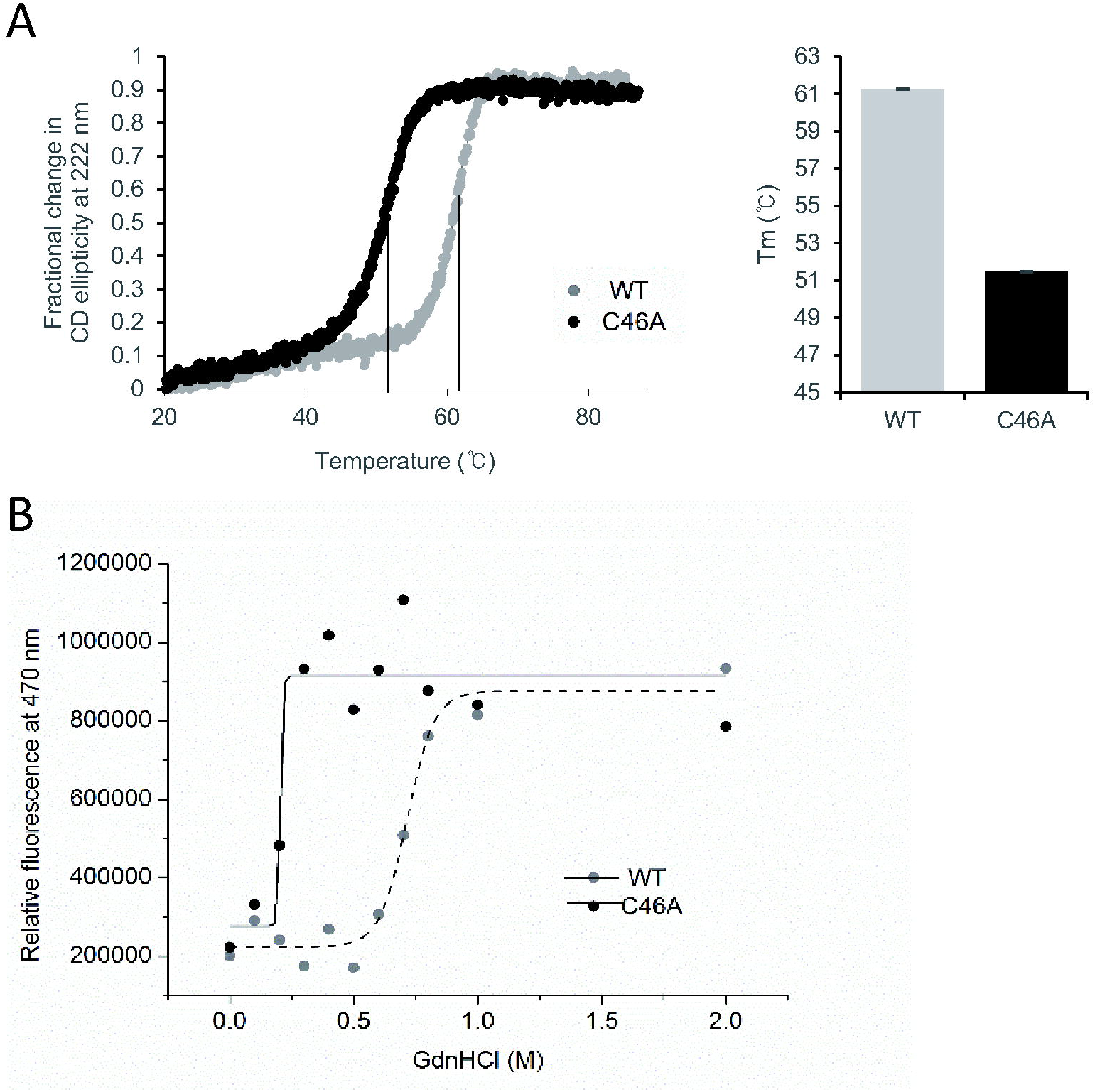
Protein stability and folding of DJ-1 proteins in response to denaturants. (A) Thermal unfolding profile of DJ-1 WT and C46A mutant by measuring CD ellipticity at 222 nm. (B) Hydrophobic exposure during GdnHCl-induced unfolding was monitored by ANS fluorescence intensity at 470 nm. WT and C46A mutant were incubated for 16 h with various concentrations of GdnHCl. Samples were prepared by mixing the protein with ANS stock solution to final molar ration of 75 : 1 (ANS : protein) and equilibrating in the dark for 30 min at RT. Fluorescent emission intensity at 470 nm was measured using an excitation wavelength 380 nm. Data for each proteins were fitted with a Boltzmann curve using Origin 8.5.

### Cysteine 46 and 53 form an intra-disulfide bond in mammalian cell

In order to verify *in vitro* results occur in mammalian cells, we examined the presence of intra-disulfide bond in HeLa cells. HeLa cells transfected with Flag-WT and Cys mutants were immunoprecipitated with Flag antibody. The enriched WT and mutant DJ-1s were shown as two bands by SDS-PAGE in non-reducing conditions, while one band in reducing condition (Fig. 6A). Additionally this result was confirmed by Western analysis using Flag antibodies (Fig. 6B). These results indicate the possibility of the presence of intra-disulfide bond inside cells. This was confirmed by identifying Cys46-Cys53 intra-disulfide bond in cellular WT DJ-1 employing tandem MS analysis combining with DBond algorithm (Fig. 6C). Also, the presence of other type of intra-disulfide bond were detected in cellular Cys mutants as well as in recombinant ones (Supplementary Table 4). Cys53-Cys106 intra-disulfide bond in C46A mutant was also identified with low intensity by tandem MS spectra (Supplementary Fig. 6). C53A mutant also showed down-shifted band to different position, suggesting that C53A mutant is presumed to form Cys46-Cys106 intra-disulfide bond. However, cellular DJ-1 did not form oligomer in response to oxidative stress.

**Fig. 6.**
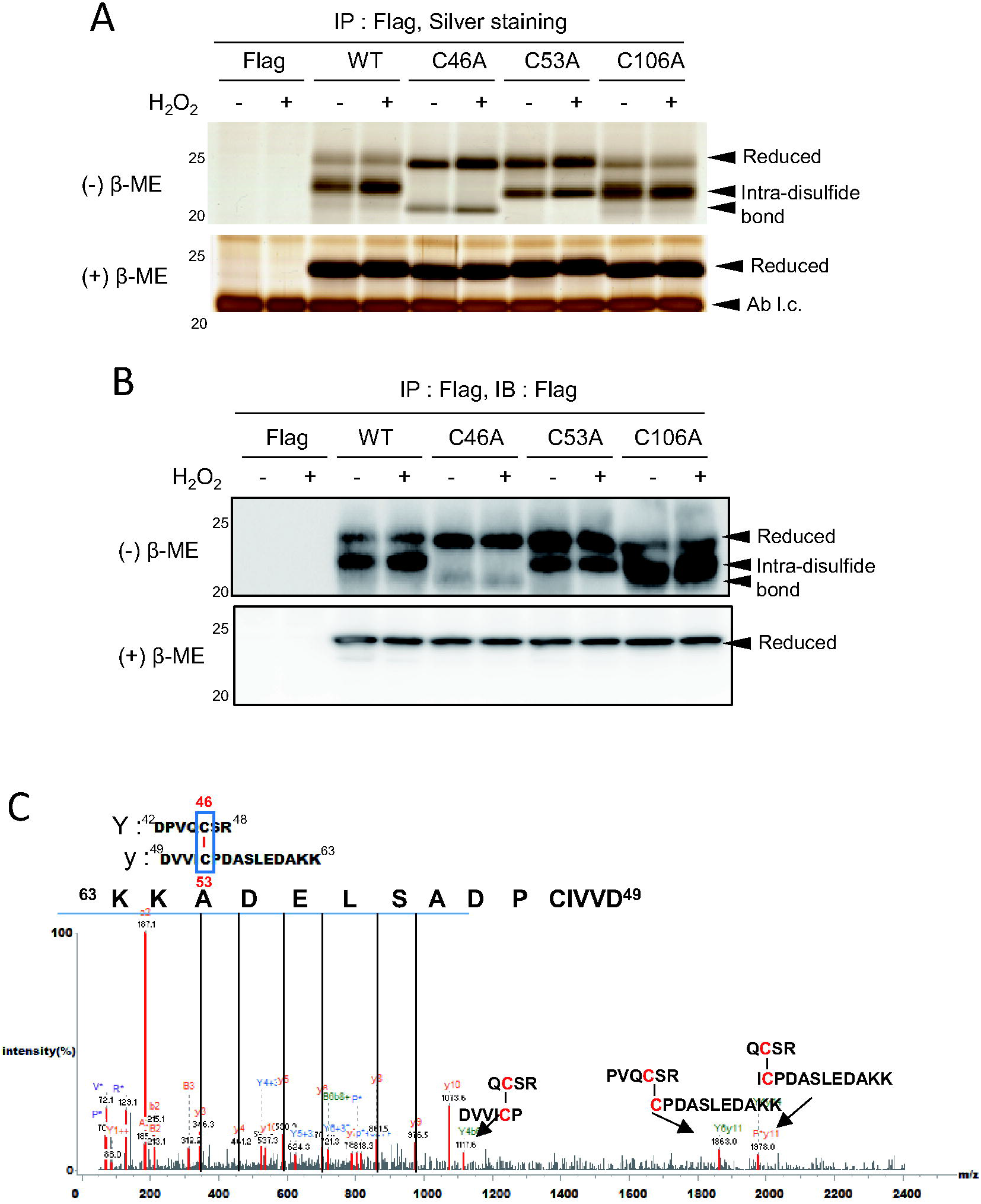
DJ-1 has Cys46-Cys53 disulfide bond in mammalian cell. (A, B) HeLa cells transfected with Flag-DJ-1 WT and Cys mutants were treated with 0 or 0.5 mM H_2_O_2_ in 37°C for 1 h. The cell lysate were subjected to immunoprecipitation (IP) with anti-Flag antibody. Immune complexes were separated by SDS-PAGE under non-reducing (−β-ME) or reducing (+β-ME) conditions and detected by (A) silver staining and (B) Western analysis using anti-Flag antibody. (C) Tandem mass spectrum of Cys46-Cys53 disulfide bond peptide obtained from WT DJ-1 without H_2_O_2_ treatment under non-reducing condition in Fig. 6A.

Oxidative modifications of cellular DJ-1s showed similar tendency with *in vitro* results with recombinant proteins, but the details were slightly different (Supplementary Fig. 7, Supplementary Table 5). Cys106s in C46A mutant were more readily oxidized to sulfonic acid than WT. Cys106s in C53A mutant were also more readily oxidized than WT to sulfinic acid, sulfonic acid and thiosulfonic acid including Cys to Ser oxidation. Cys106 in Cys mutants not forming intra-disulfide bond, is more oxidized to irreversible oxidation. The difference in detail from *in vitro* results is assumed to be due to cellular equilibrium and complexity of cellular environment including protein-protein interaction etc.

### Intra-disulfide bond formation is required for DJ-1 cellular function

Since protein structure is closely related to function, we investigated cellular functions of DJ-1 and its mutants. We examined the function of DJ-1 as a ROS scavenging protein [9] in SN4741 cells, dopaminergic neuronal cell line, and DJ-1 KO SN4741 cells derived from the *substantia nigra* of transgenic DJ-1 *null* mouse embryos in order to exclude endogenous DJ-1 activity [46, 47]. Several studies have reported that DJ-1 KO cells have increased ROS production compared to WT in the muscle [48], hematopoietic stem cell [49] and dopaminergic cell in *Drosophila* [46]. To identify whether DJ-1 deficiency is associated with cellular ROS level in dopaminergic cells as in the previous studies, we measured fluorescence generated by ROS after loading the cells with CM-H_2_DCFDA in WT and DJ-1 KO SN4741 cells. DJ-1 KO SN4741 cells showed increased cellular ROS levels (Fig. 7A). Next, we examined whether cellular ROS level increased by DJ-1 KO is lowered when DJ-1 is added back. Intracellular ROS levels were measured in DJ-1 KO SN4741 cells transfected with WT and three Cys mutants. Overexpression of WT and Cys mutants did not affect the intracellular ROS levels, however, when transfected cells were treated with 0.5 mM H_2_O_2_ for 20 min at 33°C, overexpression of WT DJ-1, not Cys mutants, decreased intracellular ROS level (Fig. 7B). Thus, the results suggest that all three Cys mutants, unlike WT DJ-1, lost their ROS scavenging activity as antioxidant protein.

**Fig. 7.**
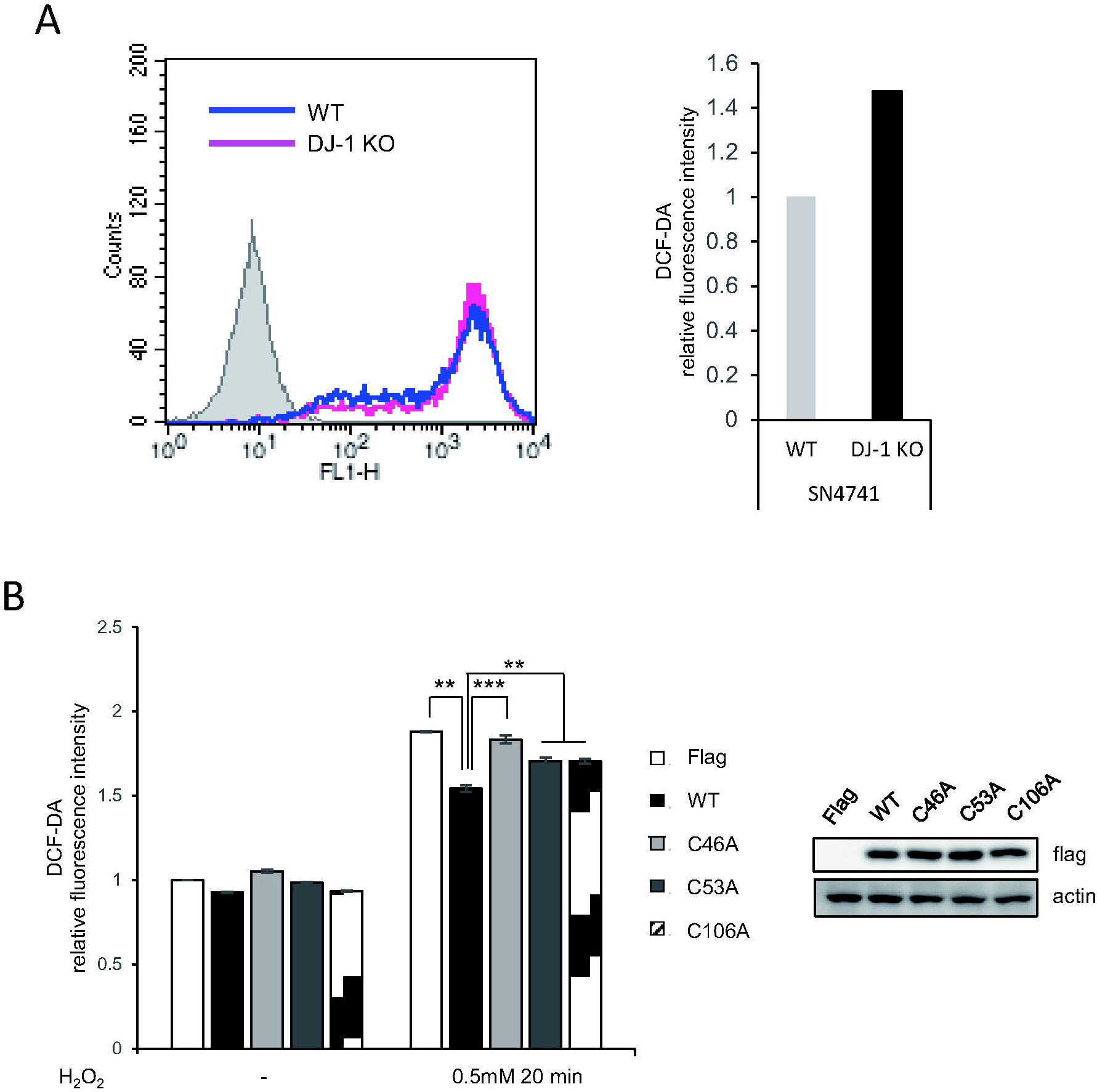
All three cysteine residues in DJ-1 are required for its role in antioxidant activity in neuronal cells. (A) Intracellular ROS level of WT and DJ-1 KO SN4741 cells were determined by FACS using H_2_DCF-DA. Equal numbers of cells were treated with 2 μM for 15 min at 33°C and immediately, the fluorescence intensity was measured. Relative ROS level is presented by Geomean value. (B) DJ-1 KO SN4741 cells transfected with Flag-DJ-1 WT and Cys mutants were treated with 0 or 0.5 mM H_2_O_2_ for 20 min at 33°C and then with H_2_DCF-DA (2 μM) for 15 min in 33°C, fluorescence intensity was measured. Data are presented as mean ± S.D. of triplicate experiments (Two tailed t-test; \*\**P* < 0.01, *** *P*<0.001).

DJ-1 is known to protect neurons against damage though mediating cell proliferation by activating the extracellular signal-regulated kinase (ERK1/2) pathway [50]. To investigate whether SN4741 cells are also enabled for DJ-1 dependent ERK signaling, WT and DJ-1 KO SN4741 cells were treated with 0.5 mM H_2_O_2_ at 33°C for indicated times and phosphorylation levels of ERK were measured using Western analysis. ERK activation was reduced in DJ-1 KO cells than WT (Fig. 8A). Then, DJ-1 KO SN4741 cells transiently transfected with Flag empty vector and Flag-DJ-1 WT were treated with 0.5 mM H_2_O_2_ at 33°C for various times. Overexpression of WT DJ-1 increased the phosphorylation level of ERK signaling, especially when H_2_O_2_ was treated for 20 min, the maximum difference in ERK activation was observed (Fig. 8B). When DJ-1 KO SN4741 cells transfected with Flag, WT and three Cys mutants were treated with 0.5 mM H_2_O_2_ for 20 min, ERK activations are shown in Fig. 8C. Overexpression of WT increased ERK activation, but all three Cys mutants did not. The results also suggest that all three Cys residues are required to activate ERK signal transduction.

**Fig. 8.**
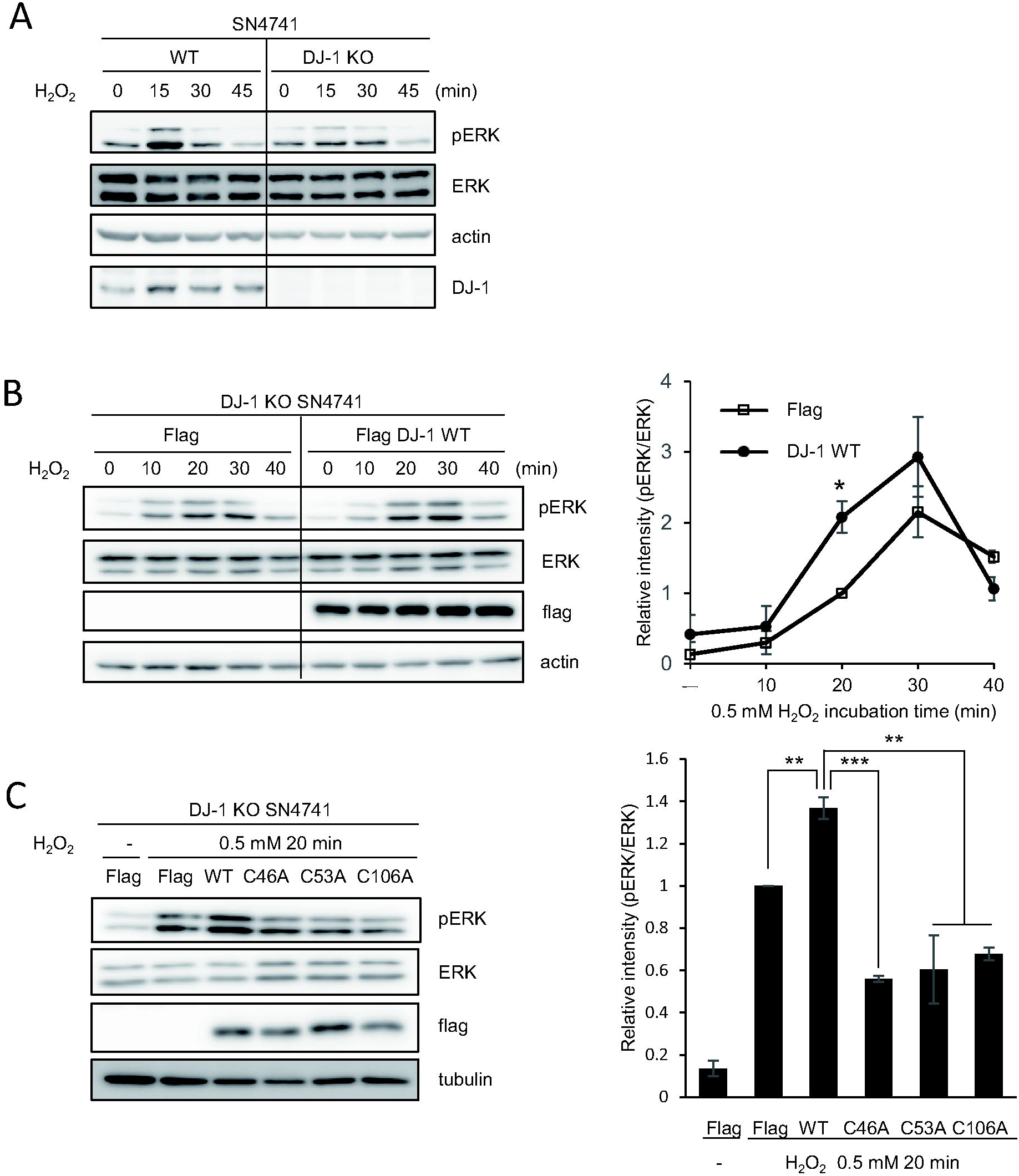
All cysteine residues in DJ-1 are required for its role in ERK activation in neuronal cells. (A) WT and DJ-1 KO SN4741 cells were treated with H_2_O_2_ (0.5 mM) at 33°C for various times (0, 15, 30 and 45 min), then cells were lysed and ERK activations were detected by Western analysis using anti-phospho-ERK(T202/Y204) antibodies. (B) DJ-1 KO SN4741 cells transiently transfected with Flag or Flag DJ-1 WT were treated with H_2_O_2_ (0.5 mM) at 33°C for various times (0, 10, 20, 30 and 40 min), then cells were lysed and ERK activations were detected by Western analysis using anti-phospho-ERK antibodies. Quantitative analysis was done with multi-gauge software. (C) DJ-1 KO SN4741 cells transiently transfected with WT and Cys mutant Flag-DJ-1s were treated with H_2_O_2_ (0.5 mM) at 33°C for 20 min and then ERK activations were detected. Data are presented as mean ± S.D. of triplicate experiments (Two tailed t-test; \**P* < 0.05, \*\**P* < 0.01 and \*\*\**P* < 0.001).

Since ERK signaling pathways in cell proliferation play a critical role in neuroprotection [51, 52], we examined cell proliferation in DJ-1 KO SN4741 cells transfected with Flag, WT and three Cys mutants employing a real-time cell analyzer, xCelligence, based on the conductivity of attached cells on the gold surface. There were no discernible differences in the proliferation of DJ-1 KO SN4741 cells transfected with WT and Cys mutants without oxidative stress (Fig. 9A). However, the apparent increase of proliferation in response to oxidative stress was detected in cells transfected with WT DJ-1, not three Cys mutants (Fig. 9B). These results demonstrate that all three Cys residues are necessary and Cys46-Cys53 intra-disulfide bond formation is required to play the cellular function of DJ-1.

**Fig. 9.**
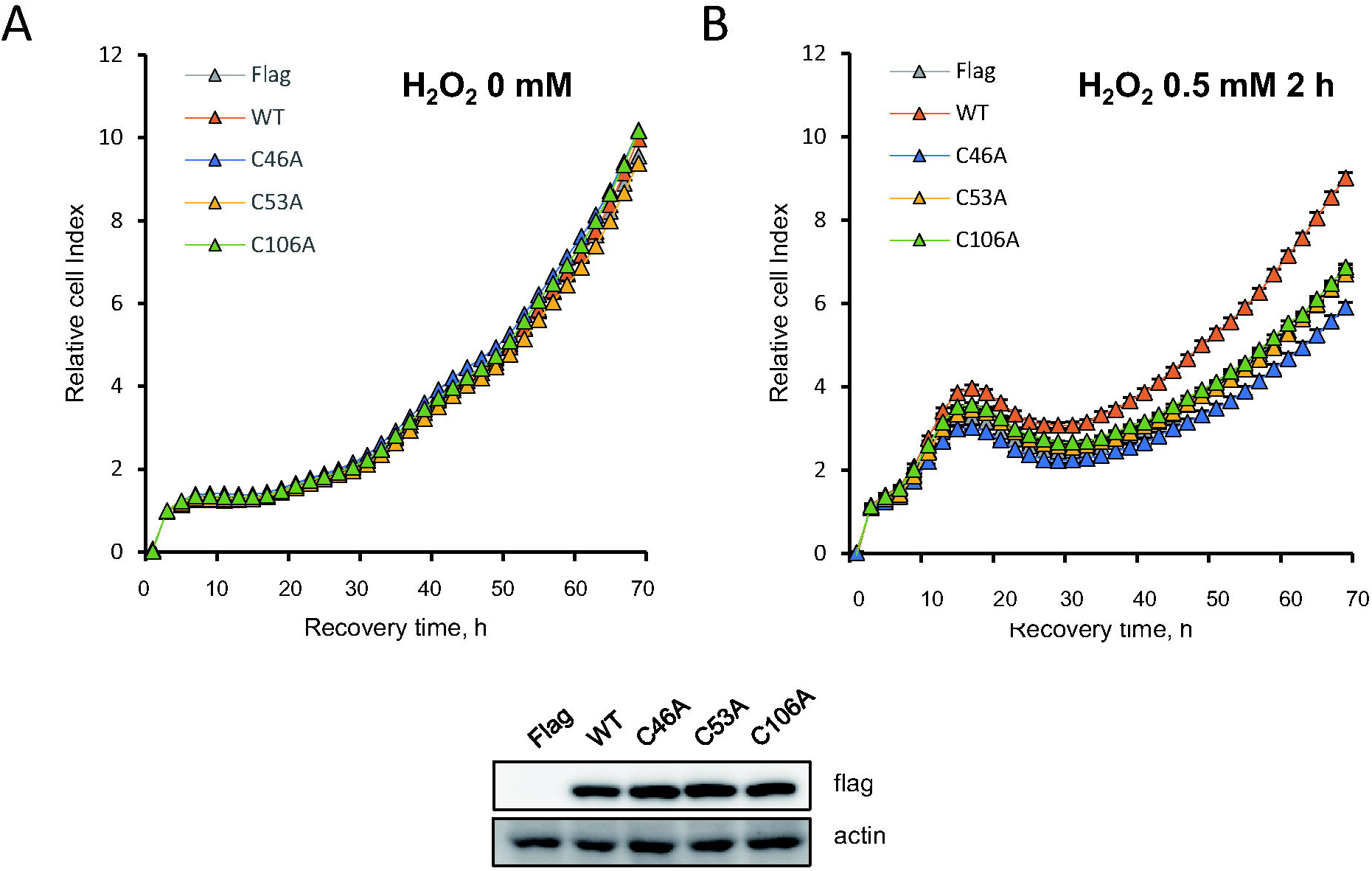
All cysteine residues of DJ-1 are essential for cell proliferation in response to oxidative stress. (A, B) SN4741 DJ-1 KO cells transiently transfected with Flag empty vector, Flag-DJ-1 and Flag-DJ-1 Cys mutants were treated with (A) 0 or (B) 0.5 mM of H_2_O_2_ at 33°C for 2 h. Cells were then plated in 96-well plate, incubated at 33°C for 3 days. Cell growth was monitored under real-time cell analyzer, xCELLigence. Data were presented as the means ± S.D. of triplicated experiments.

## Discussion

Our results establish that Cys46 in DJ-1 is also reactive cysteine and is easily oxidized to form Cys46-Cys53 intra-disulfide bonds, preventing active site Cys106 from being over-oxidized. Cys mutants, not able to form intra-disulfide bonds lose their antioxidant activity due to structural changes, which prevents them from functioning to promote cell proliferation. Thus, Cys46-Cys53 intra-disulfide bond in DJ-1 is believed to play an important role in structural integrity by maintaining the proper oxidative state of Cys106 and in antioxidant activity to protect cells from oxidative stress.

This is the first study focusing on the reactivity of Cys46 residues and the resulting Cys46-Cys53 intra-disulfide binding in the structure and function of DJ-1. Among three cysteine residues in DJ-1, studies on Cys106 have been more advanced than the other cysteine residues, and several studies have reported that Cys106 in DJ-1 is the most reactive and readily oxidized [30, 33]. Unlike Cys106, Cys46 and Cys53 are not modified but their mutants lose their function. This suggests that Cys46 and Cys53 play key roles in dimer formation via hydrophobic interaction of surface [33, 35]. However, both C46A and C53A mutants also can produce dimers *via* DSS crosslinker [35], which correlates with the results of size-exclusion chromatography in which all three Cys mutants were separated at same retention time (data not shown). The C46A mutant has the result of not being able to create a dimer in the cell [35], so the results of the dimer formation are still controversial. This suggests the possibility that inactivation of the C46A and C53A mutants may be due to disulfide bond formation, not dimer formation. Previous study has reported that DJ-1 is a redox-dependent chaperone preventing α-synuclein aggregate formation [53]. They showed the chaperone ability of DJ-1 was decreased when assay was performed in a reduced environment using reducing agent, DTT. They interpreted redox regulation governs DJ-1 chaperone activity. However, this study indicates another possibility that intra-disulfide bond of DJ-1 is necessary for functioning as a chaperone. Therefore, this study suggests for the first time that malfunction of Cys mutants can be derived from disability to form intra-disulfide bond.

Redox sensitive Cys residues can be modified to numerous ways. However, Cys46, the reactive Cys residue in DJ-1, did not show various oxidative modifications. This indicates that Cys46 has a strong tendency to form intra-disulfide bond with adjacent Cys53 rather than oxidative modifications. The consequent structural stability obtained by intra-disulfide bond maintains the reactivity of Cys106, which can be a mechanism for regulating the activity of Cys106. There is a similar example where three Cys residues work cooperatively. A tumor metastasis suppressor protein, Nm23-H1/NDPK-A, is a redox sensitive protein with three cysteine residues [54]. The most reactive cysteine, a peroxidatic cysteine, Cys4 in Nm23-H1 also easily forms an intra-disulfide bond with adjacent resolving cysteine. Therefore, oxidative modifications were not observed in Cys4 residue as Cys46 of DJ-1 [24]. Cys4 forms intra-disulfide bond with Cys145 in response to oxidative stress, thereby Cys109 residue is exposed to the surface and oxidized to sulfonic acid through a quaternary structural change, which lost the enzymatic activity of NDPK-A. Both cases show how stepwise oxidation regulates protein structure and activity. These findings broaden knowledge about stepwise molecular regulations of cysteine residues. Therefore, this study provides a new perspective for further studying and designing new experiments to reveal regulation mechanisms of DJ-1 in reactive cysteines cooperatively.

We have shown a pattern of oxidative modifications at cysteine residues of WT and mutant DJ-1s (Fig. 3). The analysis with the combined result provides conclusive behavior among cysteines. All three cysteines collaborate together for proper molecular functions. Cys46 and Cys53 form intra-disulfide bonds, but the modification of the two is different. Compared to Cys46, Cys53 has been oxidized in various forms. As already known, Cys46 is structurally embedded in the protein. So, despite being reactive by itself, no modification is observed because it forms disulfide with the nearby Cys53. On the other hand, since Cys53 is present on the protein surface, it is easy to react with a solvent, so it is thought that it is oxidized in various forms as well as intra-disulfide bonds. In addition, intra-disulfide bond between Cys46 and Cys53 is also crucial to keep the oxidation state of Cys106 not into sulfonylation but sulfinylation. Previous studies also suggest that the degree of oxidation of Cys106 is dependent on the presence or absence of Cys46 and Cys53 [55, 56]. The results suggest that disulfide formation modulates redox state of the other cysteine residue.

Since intra-disulfide bond formation is a typical modification that alters the structure of proteins, structural dynamics were examined by HDX-MS. Previous studies have reported that Cys106 is located in a putatively nucleophile elbow region, the sharp turn between a β-sheet and an α-helix, in solvent accessible branched [12, 37, 57]. Although Cys106 is known as the most reactive Cys residue and readily oxidized, it is structurally buried and could be controlled precisely. The crystal structure of native DJ-1 (Supplementary Fig. 8) implies the spatial relation of three cysteine residues and explains the possible formation of Cys53-Cys106 intra-disulfide bond in C46A mutant. In the crystal structure, Cys106 is regulated by Glu18 residing on αA helix which interacts with a flexible domain composed of four β-strands (β2-5). Intriguingly, two cysteine residues, Cys46 and Cys53, are located on a β-sheet composed of β2 and β4. However, in the case of C46A mutant, Cys53-Cys106 intra-disulfide bond was formed as proven by HDX-MS experiment. It is plausible due to the presence of two long hinge loops of flexible domain. Since there is no intra-disulfide bond in Cys46-Cys53, the flexible domain can be inverted. HDX-MS strongly supports this situation as shown in the increased deuterium exchange level in this domain. In addition, deuterium exchange levels of peptide containing Cys106 and peptides containing Glu18 and His126 were also increased. The rapid deuterium exchange at the catalytic site confirms the movement of adjacent helix αA and thus indicates the occurrence of large conformational changes in C46A mutant.

In summary, the results suggest that intra-disulfide bond and cysteine regulation of DJ-1 are important for its structure and function and broaden the understanding of the complicated regulatory mechanisms of DJ-1 that operate under oxidative environments. To understand the regulation mechanism how to be modulated between cysteine residues of DJ-1 provides insight to understand neurodegenerative disease and find a novel therapy focusing on DJ-1.

## Data availability

The structures presented in this paper have all been deposited in the Protein Data Bank (PDB) with the following codes: 4RKW

Raw data for DJ-1 PTM analysis and HDX analysis (Figure 1C, 2B, 3, 4, 6, Table 1, Supplementary Table 2-5) are available via ProteomeXchange with identifier PXD020690 [60]. Reviewer account details are as follows; Username, Password: QXG0L9LO.

## Acknowledgements

We are grateful to Jin Son (Ewha Womans University, Korea) for donating SN4741 and DJ-1(−/−) cell lines. Authors appreciate technical support of MS analysis performed by Mr. Kang W.

## Funding and additional information

This work was supported by NRF grant (No. 2020R1F1A1055369) of National Research Foundation of Korea. I.K. Song was supported by Brain Korea 21 Plus (BK21 Plus) Project.

## Author contributions

IKS and KJL designed the study, IKS conducted most of the experiments and analyzed the results. MSK and DHS designed and performed biophysical assay, and JEF did orthogonal sequence analysis. IKS, KJL and DHS wrote the paper.

