## supplementary data for "Stepwise Oxidations Play Key Roles in the Structural and Functional Regulations of DJ-1"

Table of contents

Supplementary Table 1..........S-1
Supplementary Table 2..........S-3
Supplementary Table 3..........S-4
Supplementary Table 4..........S-5
Supplementary Table 5..........S-6
Supplementary Figure 1..........S-7
Supplementary Figure 2..........S-8
Supplementary Figure 3..........S-9
Supplementary Figure 4..........S-15
Supplementary Figure 5..........S-16
Supplementary Figure 6..........S-17
Supplementary Figure 7..........S-18
Supplementary Figure 8..........S-19

**SupplementaryTable 1**. Orthologous analysis of DJ-1

| No. | Classes | Species | KEGG symbol | C46 | C53 | C106 |
|---|---|---|---|---|---|---|
| 1 | MAMMALS | Humans | hsa | C | C | C |
| 2 | | Pan troglodytes (chimpanzee) | ptr | C | C | C |
| 3 | | Pan paniscus (bonobo) | pps | C | C | C |
| 4 | | Gorilla gorilla gorilla (western lowland gorilla) | ggo | C | C | C |
| 5 | | Pongo abelii (Sumatran orangutan) | pon | C | C | C |
| 6 | | Nomascus leucogenys (northern white-cheeked gibb | nle | C | C | C |
| 7 | | Macaca mulatta (rhesus monkey) | mcc | C | C | C |
| 8 | | Macaca fascicularis (crab-eating macaque) | mcf | C | C | C |
| 9 | | Callithrix jacchus (white-tufted-ear marmoset) | cjc | C | C | C |
| 10 | | Mus musculus (mouse) | mmu | C | C | C |
| 11 | | Rattus norvegicus (rat) | rno | C | C | C |
| 12 | | Cricetulus griseus (Chinese hamster) | cge | C | C | C |
| 13 | | Nannospalax galili (Upper Galilee mountains blind mo | ngi | C | C | C |
| 14 | | Heterocephalus glaber (naked mole rat) | hgl | C | C | C |
| 15 | | Oryctolagus cuniculus (rabbit) | ocu | C | C | C |
| 16 | | Tupaia chinensis (Chinese tree shrew) | tup | C | C | C |
| 17 | | Canis familiaris (dog) | cfa | C | C | C |
| 18 | | Ailuropoda melanoleuca (giant panda) | aml | C | C | C |
| 19 | | Ursus maritimus (polar bear) | umr | C | C | C |
| 20 | | Felis catus (domestic cat) | fca | C | C | C |
| 21 | | Panthera tigris altaica (Amur tiger) | ptg | C | C | C |
| 22 | | Bos taurus (cow) | bta | C | C | C |
| 23 | | Bos mutus (wild yak) | bom | C | C | C |
| 24 | | Pantholops hodgsonii (chiru) | phd | C | C | C |
| 25 | | Capra hircus (goat) | chx | C | C | C |
| 26 | | Ovis aries (sheep) | oas | C | C | C |
| 27 | | Sus scrofa (pig) | ssc | C | C | C |
| 28 | | Camelus ferus (Wild Bactrian camel) | cfr | C | C | C |
| 29 | | Balaenoptera acutorostrata scammoni (minke whale) | bacu | C | C | C |
| 30 | | Lipotes vexillifer (Yangtze River dolphin) | lve | C | C | C |
| 31 | | Equus caballus (horse) | ecb | C | C | C |
| 32 | | Myotis brandtii (Brandt's bat) | myb | C | C | C |
| 33 | | Myotis davidii | myd | C | C | C |
| 34 | | Pteropus alecto (black flying fox) | pale | C | C | C |
| 35 | | Monodelphis domestica (opossum) | mdo | C | C | C |
| 36 | | Sarcophilus harrisii (Tasmanian devil) | shr | C | C | C |
| 37 | | Ornithorhynchus anatinus (platypus) | oaa | C | C | C |
| 38 | BIRDS | Gallus gallus (chicken) | gga | C | C | C |
| 39 | | Meleagris gallopavo (turkey) | mgp | C | C | C |
| 40 | | Anas platyrhynchos (mallard) | apla | C | C | C |
| 41 | | Taeniopygia guttata (zebra finch) | tgu | C | C | C |
| 42 | | Geospiza fortis (medium ground-finch) | gfr | C | C | C |
| 43 | | Ficedula albicollis (collared flycatcher) | fab | C | C | C |
| 44 | | Pseudopodoces humilis (Tibetan ground-tit) | phi | C | C | C |
| 45 | | Corvus cornix (hooded crow) | ccw | C | C | C |
| 46 | | Falco peregrinus (peregrine falcon) | fpg | C | C | C |
| 47 | | Falco cherrug (Saker falcon) | fch | C | C | C |
| 48 | | Columba livia (rock pigeon) | clv | C | C | C |
| 49 | REPTILES | Alligator sinensis (Chinese alligator) | asn | C | C | C |
| 50 | | Alligator mississippiensis (American alligator) | amj | C | C | C |
| 51 | | Pelodiscus sinensis (Chinese soft-shelled turtle) | pss | C | C | C |
| 52 | | Chelonia mydas (green sea turtle) | cmy | C | C | C |
| 53 | | Anolis carolinensis (green anole) | acs | C | C | C |
| 54 | | Python bivittatus (Burmese python) | pbi | C | C | C |
| 55 | AMPHIBIANS | Xenopus laevis (African clawed frog) | xla | C | C | C |
| 56 | | Xenopus tropicalis (western clawed frog) | xtr | C | C | C |
| 57 | FISHES | Danio rerio (zebrafish) | dre | C | C | C |
| 58 | | Takifugu rubripes (torafugu) | tru | C | C | C |
| 59 | | Maylandia zebra (zebra mbuna) | mze | C | C | C |
| 60 | | Oryzias latipes (Japanese medaka) | ola | C | C | C |
| 61 | | Xiphophorus maculatus (southern platyfish) | xma | C | C | C |
| 62 | | Latimeria chalumnae (coelacanth) | lcm | C | C | C |
| 63 | CARTILAGINOUS FISHES | Callorhinchus milii (elephant shark) | cmk | C | V | C |
| 64 | LANCELETS | Branchiostoma floridae (Florida lancelet) | bfo | C | C | 2 C |
| 65 | ASCIDIANS | Ciona intestinalis (sea squirt) | cin | C | Q | C |
| 66 | ECHINODERMS | Strongylocentrotus purpuratus (purple sea urchin) | spu | C | V | – |

| No. | Classes | Species | KEGG symbol | C46 | C53 | C106 |
|---|---|---|---|---|---|---|
| 67 | INSECTS | Drosophila melanogaster (fruit fly) | dme | C | V | C |
| 68 | | Drosophila pseudoobscura pseudoobscura | dpo | C | V | C |
| 69 | | Drosophila ananassae | dan | C | V | C |
| 70 | | Drosophila erecta | der | C | V | C |
| 71 | | Drosophila persimilis | dpe | C | V | C |
| 72 | | Drosophila sechellia | dse | C | V | C |
| 73 | | Drosophila simulans | dsi | C | V | C |
| 74 | | Drosophila willistoni | dwi | G | V | C |
| 75 | | Drosophila yakuba | dya | C | V | C |
| 76 | | Drosophila grimshawi | dgr | G | V | C |
| 77 | | Drosophila mojavensis | dmo | C | V | C |
| 78 | | Drosophila virilis | dvi | C | V | C |
| 79 | | Musca domestica (house fly) | mde | C | V | C |
| 80 | | Anopheles gambiae (mosquito) | aga | C | K | C |
| 81 | | Aedes aegypti (yellow fever mosquito) | aag | C | K | C |
| 82 | | Culex quinquefasciatus (southern house mosquito) | cqu | – | I | C |
| 83 | | Apis mellifera (honey bee) | ame | C | H | C |
| 84 | | Solenopsis invicta (red fire ant) | soc | C | C | C |
| 85 | | Acromyrmex echinatior (Panamanian leafcutter ant) | aec | C | C | C |
| 86 | | Harpegnathos saltator (Jerdon's jumping ant) | hst | C | C | C |
| 87 | | Camponotus floridanus (Florida carpenter ant) | cfo | C | C | C |
| 88 | | Nasonia vitripennis (jewel wasp) | nvi | C | C | C |
| 89 | | Tribolium castaneum (red flour beetle) | tca | C | K | C |
| 90 | | Bombyx mori (domestic silkworm) | bmor | C | V | C |
| 91 | | Plutella xylostella (diamondback moth) | pxy | C | V | C |
| 92 | | Acyrthosiphon pisum (pea aphid) | api | T | K | C |
| 93 | | Pediculus humanus corporis (human body louse) | phu | C | M | C |
| 94 | MITES AND TICKS | Ixodes scapularis (black-legged tick) | isc | – | – | C |
| 95 | NEMATODES | Caenorhabditis elegans (nematode) | cel | C | V | C |
| 96 | | Caenorhabditis briggsae | cbr | C | V | C |
| 97 | | Brugia malayi (filaria) | bmy | C | T | C |
| 98 | | Loa loa (eye worm) | loa | C | M | C |
| 99 | | Trichinella spiralis | tsp | C | K | C |
| 100 | ANNELIDS | Helobdella robusta | hro | C | K | C |
| 101 | MOLLUSKS | Lottia gigantea (owl limpet) | lgi | C | V | C |
| 102 | | Crassostrea gigas (Pacific oyster) | crg | C | V | C |
| 103 | FLATWORMS | Schistosoma mansoni | smm | G | K | C |
| 104 | CNIDARIANS | Nematostella vectensis (sea anemone) | nve | C | K | C |
| 105 | | Hydra vulgaris | hmg | C | Q | C |
| 106 | PLACOZOANS | Trichoplax adhaerens | tad | NG | NG | NG |
| 107 | PORIFERANS | Amphimedon queenslandica (sponge) | aqu | C | V | C |

| | DJ-1 | | |
|---|---|---|---|
| | C46 | C53 | C106 |
| %Cys over organisms with the gene | 94.3 | 56.7 | 99.0566 |
| %Cys over all 107 organisms | 93.5 | 63.6 | 98.13084 |
| QUALITITATIVE CONSERVATION | HIGH | MED | HIGH |

Abbreviatons:
"–" means there is a gap at that position in the alignment.
"NG" means there is no true ortholog (no gene) in that species

**Supplementary Table 2**. Identified peptide of intra-disulfide bonds in the C46A mutant recombinant protein bands (0.1mM $H_2O_2$) under the non-reducing condition in Fig. 2a. Band was cut out, destained and in-gel digested by Glu-C and trypsin, and identified using nanoUPLC—ESI-q-TOF tandem MS.

| | Disulfide bond | Mass(m/z) Experimental (Charge state) | Mass Theoretical | Δm (Da) | Dbond Score | Sequence |
|---|---|---|---|---|---|---|
| C46A | Cys53-Cys106 | 1196.1418(2+) | 2390.2701 | -0.0011 | 2.2 | VVIC*PD-GLIAAIC*AGPTALLAHE |

**Supplementary Table 3.** Differential deuterium exchange rates of identified DJ-1 peptides in HDX-MS experiments.

| Sequence | Start | End | Domain | Structure | Differential deuterium exchange rate (%) | | | | | Mean |
|---|---|---|---|---|---|---|---|---|---|---|
| | | | | | 10 sec | 60 sec | 300 sec | 1800 sec | 10800 sec | |
| MASKRALVIL | 1 | 10 | | β1 | 20% | 30% | 20% | 20% | 20% | 22% |
| LVILAKGAEE | 7 | 16 | | | 0% | 0% | 0% | 10% | 10% | 4% |
| LAKGAEE | 10 | 16 | | | 0% | 0% | 0% | 15% | 14% | 6% |
| METVIPVD | 17 | 24 | E18 catalytic H-bond | αA | 25% | 25% | 25% | 38% | 44% | 31% |
| METVIPVDVM | 17 | 26 | | | 38% | 38% | 50% | 50% | 63% | 48% |
| RRAGIKVTVAGLAGKDPVQ | 27 | 45 | | αA β2 | 21% | 15% | 15% | 26% | 16% | 19% |
| GLAGKDPVQCSRD | 37 | 49 | C46 | β2 β3 | 29% | 37% | 19% | 19% | 10% | 23% |
| VVICPDASL | 50 | 58 | C53 | β4 β5 | 0% | 6% | 0% | 11% | 28% | 9% |
| EDAKKEGPYD | 59 | 68 | | β6 αB | 20% | 15% | 10% | 0% | 0% | 9% |
| EDAKKEGPYDVVVLPGGNLGAQNL | 59 | 82 | | β6 αB β6 | 9% | 13% | 4% | 8% | 4% | 8% |
| ED AKKEGPYDVVVLPGGNLGAQNLSESAAVKEIL | 59 | 92 | | | 5% | 14% | 14% | 17% | 21% | 14% |
| PGGNLGAQNL | 73 | 82 | | β6 | 10% | 10% | 0% | 0% | 5% | 5% |
| SESAAVKEIL | 83 | 92 | | αC | 0% | 10% | 0% | 10% | 10% | 6% |
| SESAAVKEILKEQENRKGLIA | 83 | 103 | | αC αD | 24% | 29% | 19% | 19% | 9% | 20% |
| KEQENRKGLIA | 93 | 103 | | αD | 9% | 9% | 18% | 9% | 13% | 12% |
| LIAAICAGPT | 101 | 110 | C106 catalytic site | β7 | 27% | 27% | 27% | 27% | 18% | 25% |
| AICAGPTAL | 104 | 112 | | β7 αE | 6% | 11% | 11% | 27% | 16% | 14% |
| AICAGPTALL | 104 | 113 | | β7 αE | 6% | 11% | 11% | 27% | 16% | 14% |
| TALLAHEIGF | 110 | 119 | | αE | 10% | 5% | 0% | 0% | 0% | 3% |
| IGFGSKVTTH | 117 | 126 | H126 catalytic diad | β8 | 10% | 20% | 20% | 30% | 35% | 23% |
| IGFGSKVTTHPLAKDKM | 117 | 131 | | β8 | 20% | 20% | 7% | 7% | 7% | 12% |
| G SKVTTHPLAKDKMMNGGHYTYSEN | 120 | 144 | | β8 αF | 0% | 4% | 0% | 4% | 0% | 2% |
| VTTHPLAKDKMMN | 123 | 135 | | β8 αF | 0% | 0% | 8% | 0% | 0% | 2% |
| MNGGHYTYSEN | 132 | 144 | | αF β9 | -8% | -16% | -8% | -15% | -15% | -12% |
| NRVEKDGL | 144 | 151 | | β9 β10 | 7% | 0% | 12% | 12% | 25% | 11% |
| RVEKDGLIL | 145 | 153 | | β10 β11 | 11% | 11% | 5% | 11% | 11% | 10% |
| ILTSRGPGTSFEF | 152 | 163 | | β11 | 9% | 4% | 0% | 0% | 0% | 3% |
| FALAIVE | 164 | 170 | Helix-kink-helix motif | αG | 14% | 29% | 36% | 43% | 50% | 34% |
| AIVEALNGKEVAAQVKAPLVL | 167 | 187 | | αG αH | 19% | 14% | 4% | 5% | 5% | 9% |
| ALNGKEVAAQVK | 171 | 182 | | αG αH | 8% | 0% | 0% | 0% | 0% | 2% |
| ALNGKEVAAQVKAPLVL | 171 | 187 | | αH | 12% | 18% | 6% | 6% | 0% | 8% |
| ALNGKEVAAQVKAPLVLKD | 171 | 189 | | αH | 16% | 16% | 0% | 0% | 0% | 6% |
| VKAPLVLKD | 181 | 189 | | αH | 0% | 7% | 0% | 0% | -7% | 0% |

- -20 ~ -11 (%)
- -10 ~ 10 (%)
- 11 ~ 20 (%)
- 21 ~ 30 (%)
- 31 ~ 40 (%)
- 41 ~ 50 (%)
- 51 ~ (%)

**Supplementary Table 4.** List of identified intra-disulfide bonds in cellular WT and C46A mutant proteins under the non-reducing condition. Immunoprecipitated proteins without oxidative stress were treated with NEM, and then analyzed with MS combining with DBond algorithm. MS/MS spectra are presented in Fig. 6c and Supplementary Fig. 2.

| | Disulfide bond | Mass(m/z) Experimental (Charge state) | Mass Theoretical | Δm (Da) | Dbond Score | Sequence |
|---|---|---|---|---|---|---|
| WT | Cys46-Cys53 | 601.7873(4+) | 2403.141 | -0.0209 | 31.1 | DPVQC*SR-DVVIC*PDASLEDAKK |
| C46A | Cys53-Cys106 | 797.7573(3+) | 2390.2701 | -0.02 | 2.8 | VVIC*PD-GLIAAIC*AGPTALLAHE |

**Supplementary Table 5**. Cysteine oxidative modifications of DJ-1 WT and mutants in Hela cells. Peptides with highest ion score were tabulated. MS/MS spectra of modified peptides are presented in Supplementary Fig. 3.

| Residue | Modification | Start-End | Mass(m/z) Experimental | Mass Theoretical | Δm (Da) | WT | C46A | C53A | C106 | Sequence |
|---|---|---|---|---|---|---|---|---|---|---|
| C106 | Cys-SH | 99-122 | 780.1008(3+) | 2337.2878 | -0.0073 | ND | 13 | 23 | ND | KGLIAAI**C**AGPTALLAHEIGFGSK |
| | Cys-$SO_2$H | 100-122 | 748.0653(3+) | 2241.1827 | -0.0083 | 32 | 41 | 83 | ND | GLIAAI**C**AGPTALLAHEIGFGSK |
| | Cys-$SO_3$H | 100-122 | 1129.5917(2+) | 2257.1776 | -0.0088 | 63 | 86 | 63 | ND | |
| | Cys to Ser | 100-122 | 732.0755(3+) | 2193.2157 | -0.0111 | ND | ND | 18 | ND | |
| | Cys-$SO_2$-SH | 100-122 | 758.7240(2+) | 2273.1548 | -0.0046 | 36 | 62 | 62 | DN | |
| C46 | Cys-SH | 33-48 | 800.9263(2+) | 1599.8403 | -0.0034 | 105 | ND | 83 | 72 | VTVAGLAGKDPVQ**C**SR |
| | Cys-$SO_3$H | 33-48 | 824.9172(2+) | 1647.8250 | -0.0052 | 68 | ND | 63 | ND | |
| | Cys to Ser | 33-48 | 528.9614(3+) | 1583.8631 | -0.0008 | 32 | ND | ND | ND | |
| | Cys-$SO_2$-SH | 33-48 | 832.9068(2+) | 1663.8022 | -0.0031 | 28 | ND | 51 | ND | |
| C53 | Cys-SH | 49-63 | 801.9047(2+) | 1601.7948 | -0.0022 | 70 | 68 | ND | 65 | DVVI**C**PDASLEDAKK |
| | Cys-$SO_2$H | 49-63 | 545.6025(3+) | 1633.7869 | -0.0012 | 30 | ND | ND | ND | |
| | Cys-$SO_3$H | 49-63 | 825.8970(2+) | 1649.7818 | -0.0024 | 58 | 62 | ND | ND | |
| | Cys to Ser | 49-63 | 529.6145(3+) | 1585.8199 | 0.0018 | 13 | ND | ND | ND | |
| | Cys-$SO_2$-SH | 49-63 | 833.8853(2+) | 1665.7590 | -0.0029 | 37 | 48 | ND | ND | |

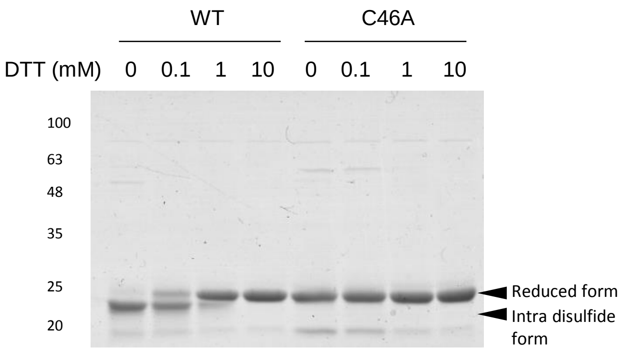


**Supplementary Figure 1.** Intra-disulfide patterns of DJ-1 WT and C46A mutant after DTT treatment. Proteins (0.5 µg) were incubated with various concentration of DTT solutions for 20 min at RT. The reactions were stopped by adding gel sample buffer and heated, and then samples were separated with non-reducing SDS-PAGE and stained with Coomassie blue.

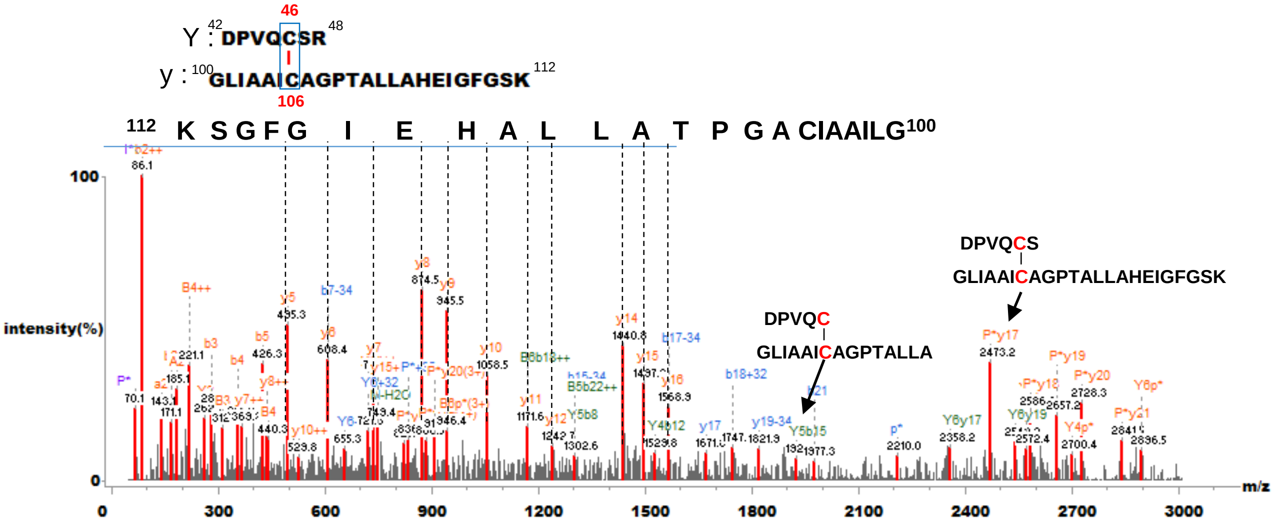


**Supplementary Figure 2**. Tandem mass spectra of Cys46-Cys106 disulfide bond peptide of the recombinant C53A mutant without $H_2O_2$ treatment in Fig. 2a Commassie-stained gels were cut out the gel and analyzed by MS and DBond analysis was performed..

$^{99}$KGIAAICAGPTALLAHEIGFGSK$^{122}$ (C106)

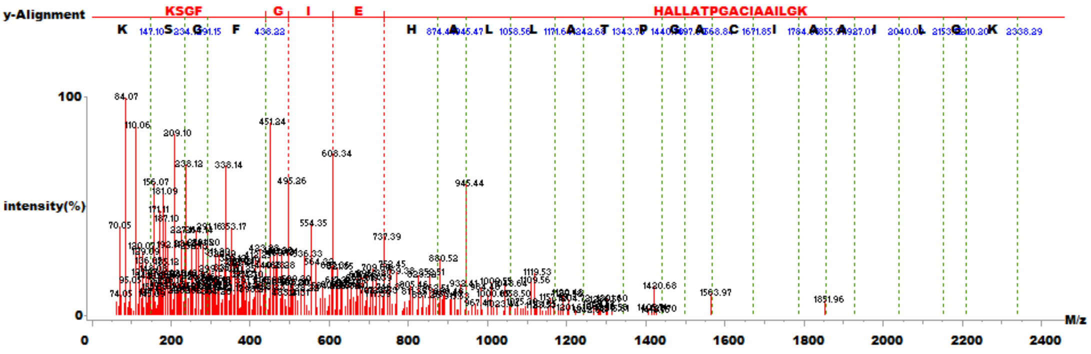


$^{100}$GIAAICAGPTALLAHEIGFGSK$^{122}$ + Cys-$SO_2H$ (C106)

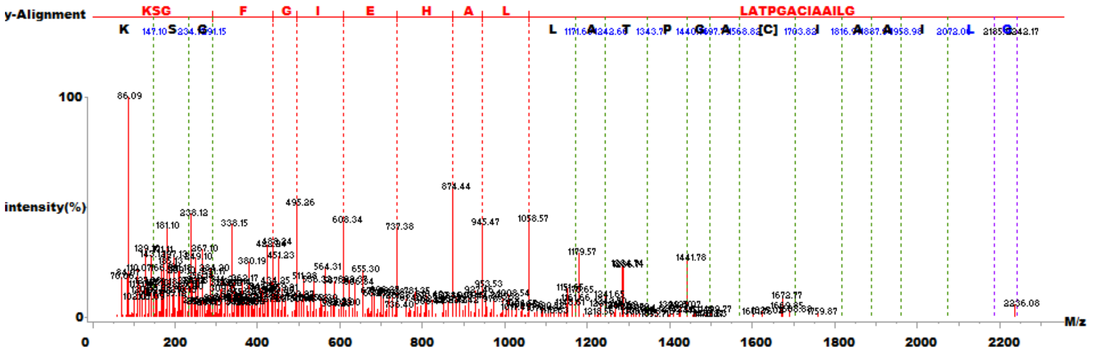


$^{99}$KGIAAICAGPTALLAHEIGFGSK$^{122}$ + Cys-$SO_3H$ (C106)

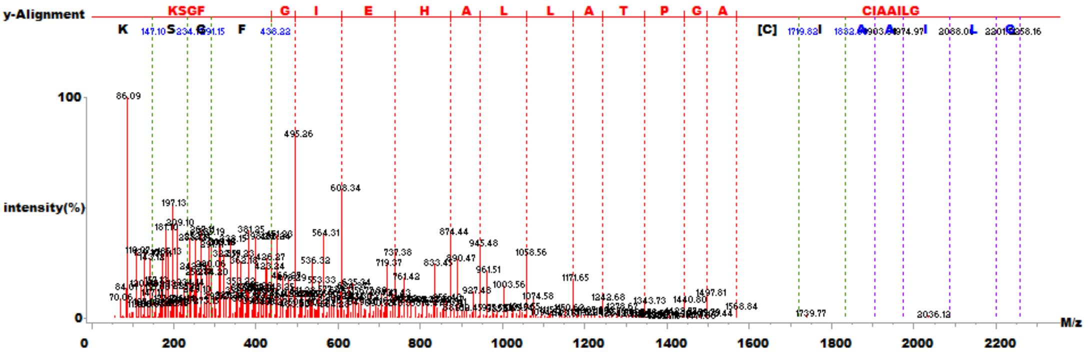


$^{100}$GIAAICAGPTALLAHEIGFGSK$^{122}$ + Cys-$SO_2$-SH (C106)

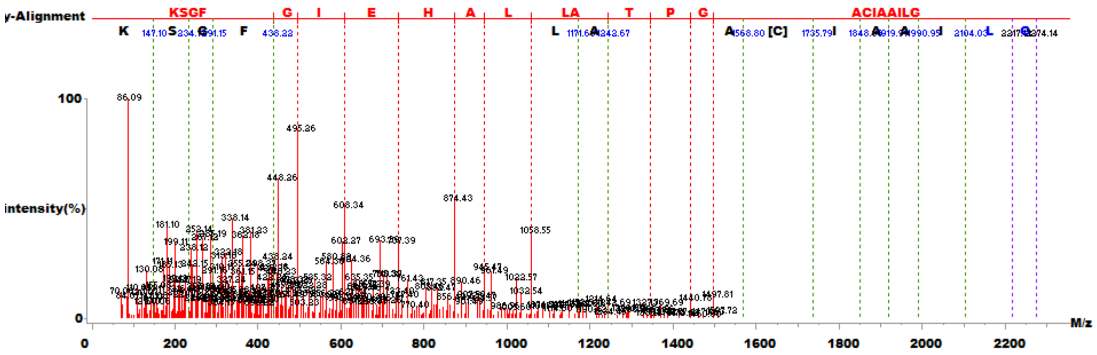


**Supplementary Figure 3.** Representative MS/MS spectra of post-translationally modified of DJ-1 listed in Fig. 3a and Supplementary Table 5 (in the order of appearance in the table).

$^{33}$VTVAGLAGKDPVQCSR$^{48}$ (C46)

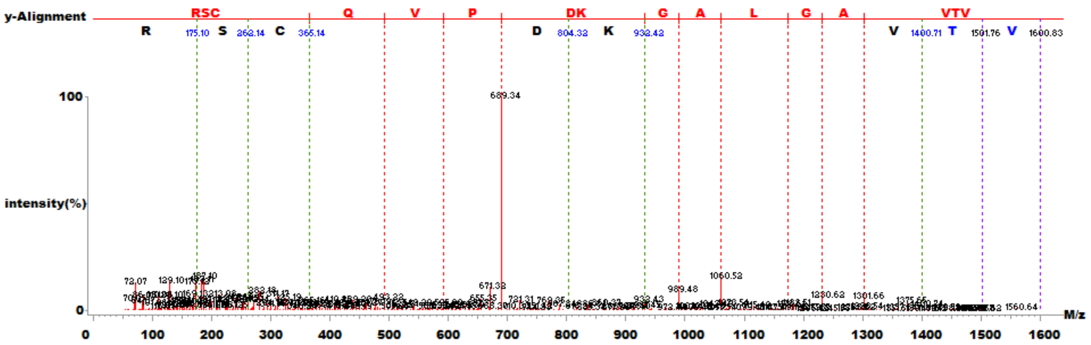


$^{33}$VTVAGLAGKDPVQCSR$^{48}$ + Cys-$SO_3H$ (C46)

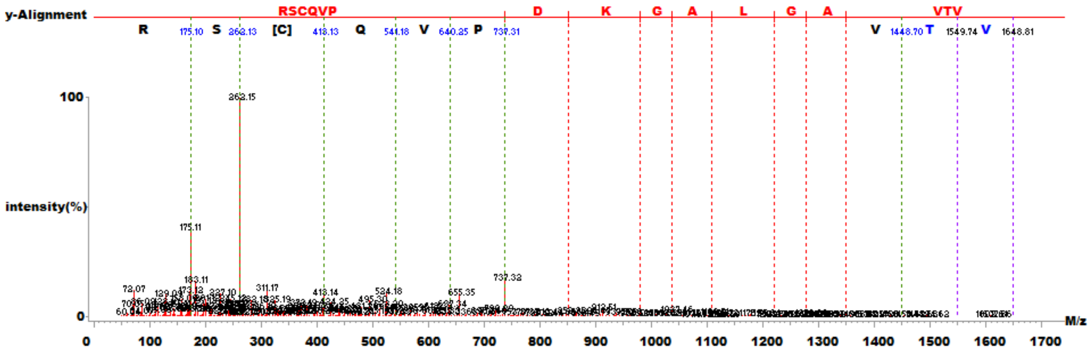


$^{49}$DVVICPDASLEDAKK$^{63}$ + Propionamide (C53)

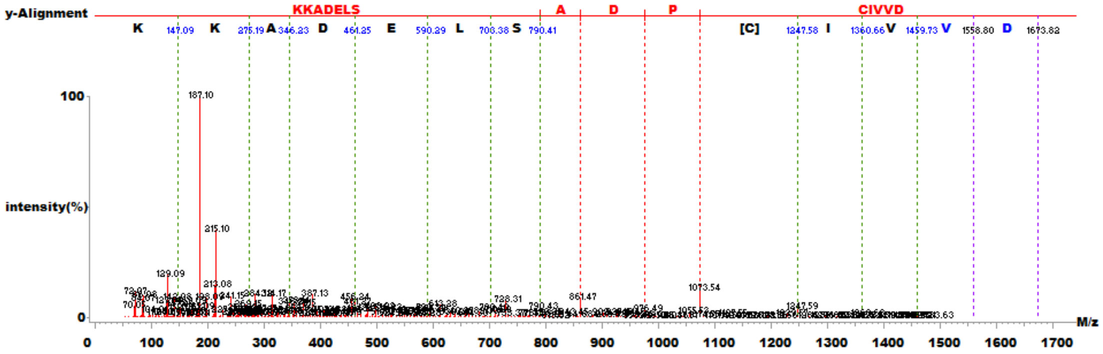


$^{49}$DVVICPDASLEDAKK$^{63}$ + Cys-$SO_2H$ (C53)

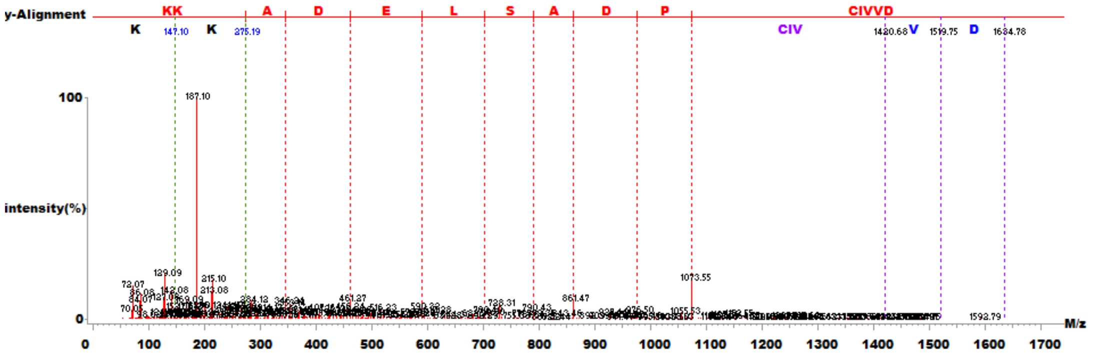


**Supplementary Figure 3.** (Continued)

$^{49}$DVVICPDASLEDAKK$^{63}$ + Cys-$SO_3H$ (C53)

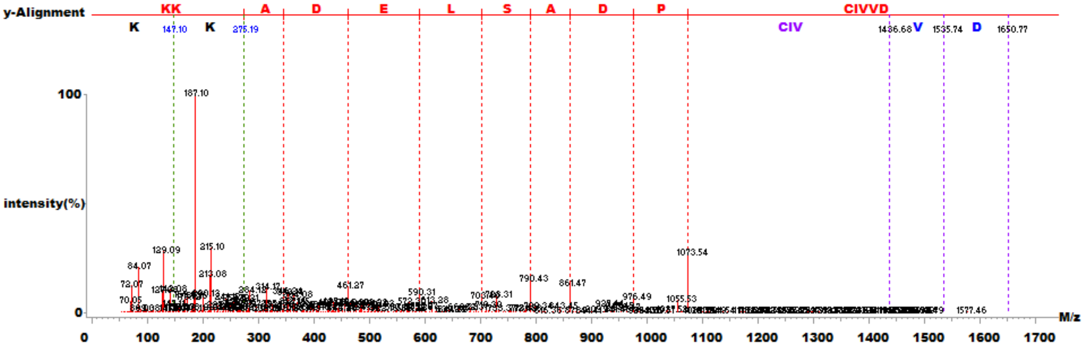


$^{49}$DVVICPDASLEDAKK$^{63}$ + Cys-$SO_2$-SH (C53)

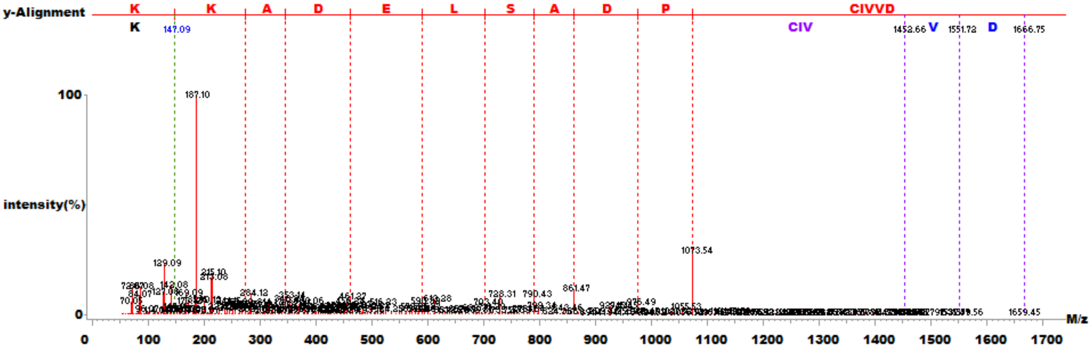


$^{99}$KGIAAICAGPTALLAHEIGFGSK$^{122}$ (C106)

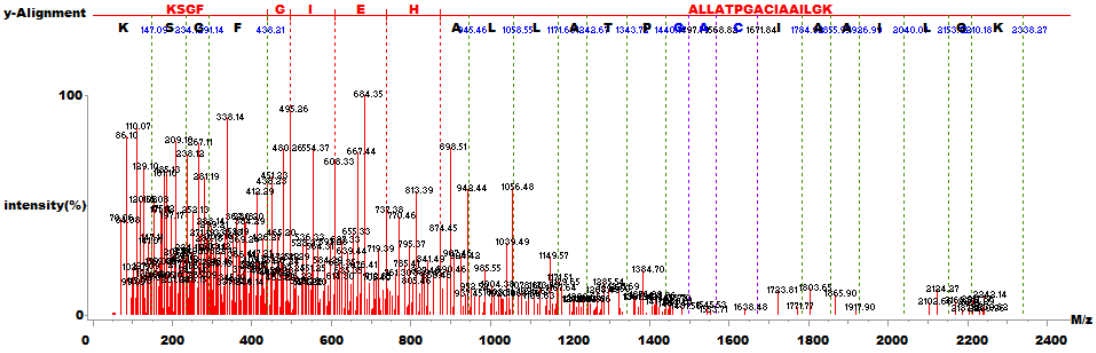


$^{100}$GIAAICAGPTALLAHEIGFGSK$^{122}$ + Cys-$SO_2H$ (C106)

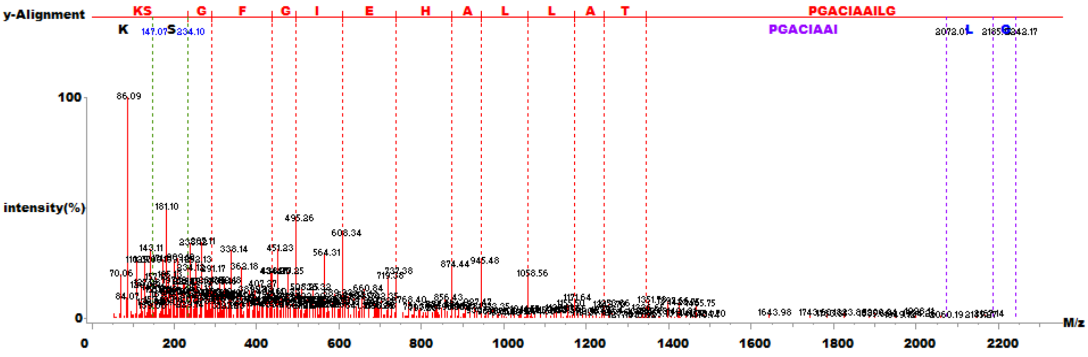


**Supplementary Figure 3.** (Continued)

$^{100}$GIAAICAGPTALLAHEIGFGSK$^{122}$ + Cys-$SO_3H$ (C106)

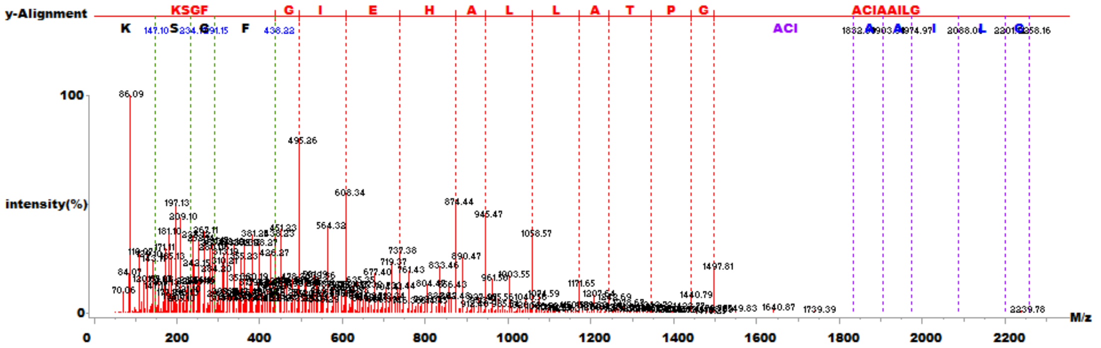


$^{100}$GIAAICAGPTALLAHEIGFGSK$^{122}$ + Cys-$SO_2$-SH (C106)

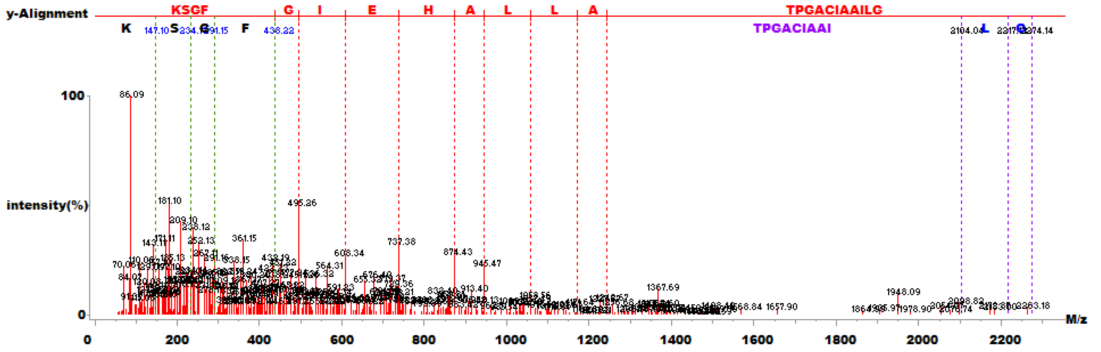


$^{100}$GIAAICAGPTALLAHEIGFGSK$^{122}$ + Cys to Ser (C106)

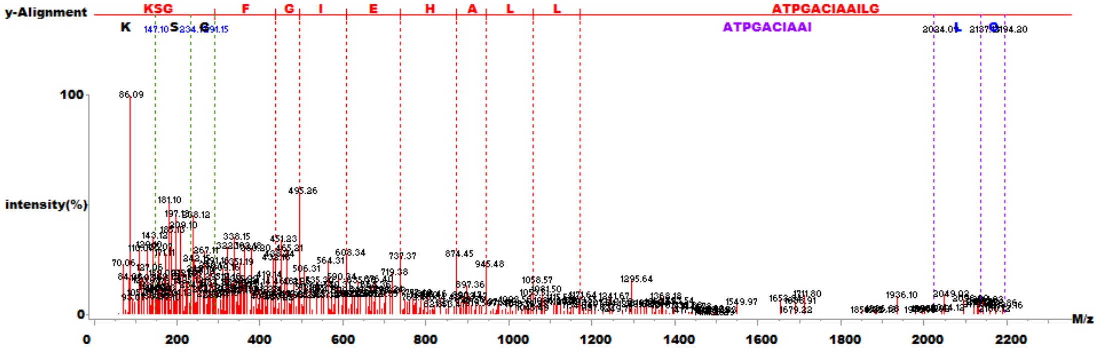


$^{33}$VTVAGLAGKDPVQCSR$^{48}$ (C46)

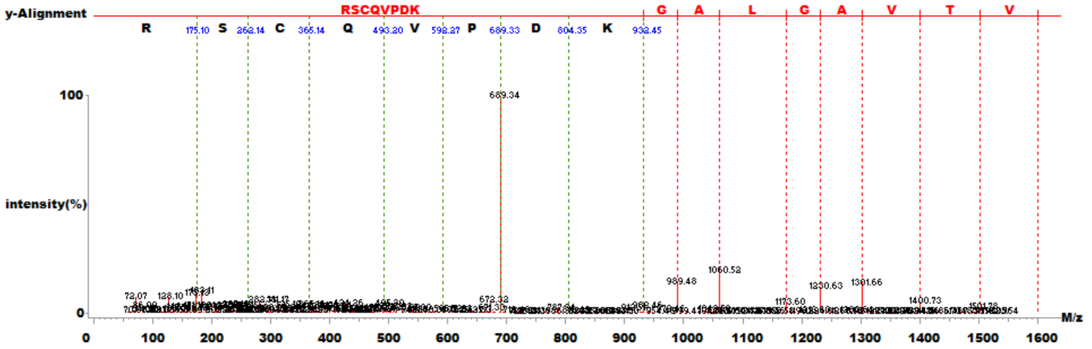


**Supplementary Figure 3.** (Continued)

$^{33}$VTVAGLAGKDPVQCSR$^{48}$ + Cys-$SO_3H$ (C46)

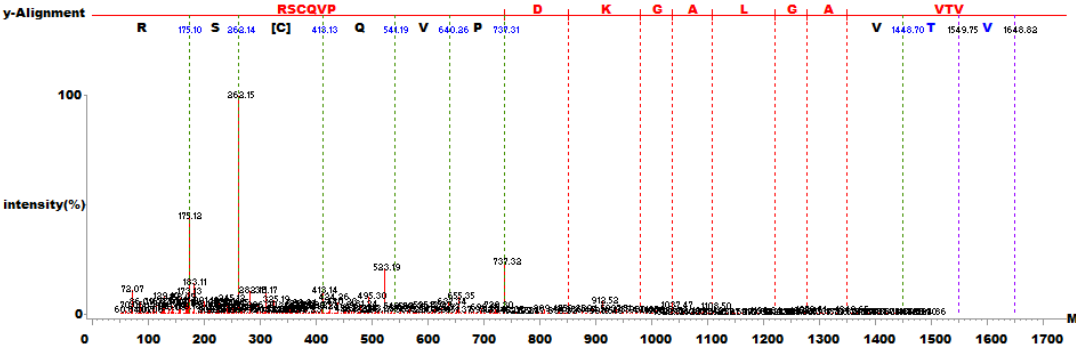


$^{33}$VTVAGLAGKDPVQCSR$^{48}$ + Cys to Ser (C46)

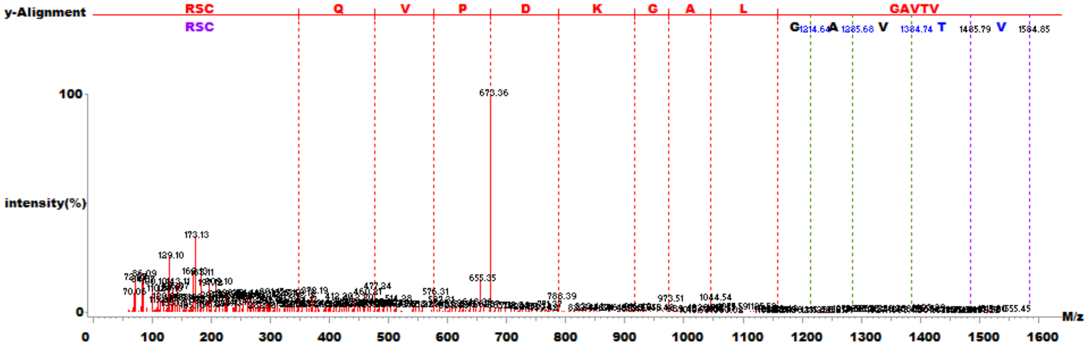


$^{33}$VTVAGLAGKDPVQCSR$^{48}$ + Cys-$SO_2$-SH (C46)

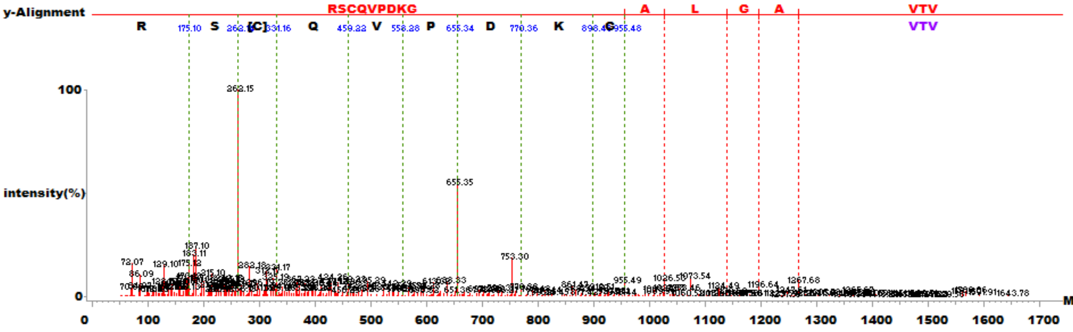


$^{49}$DVVICPDASLEDAKK$^{63}$ (C53)

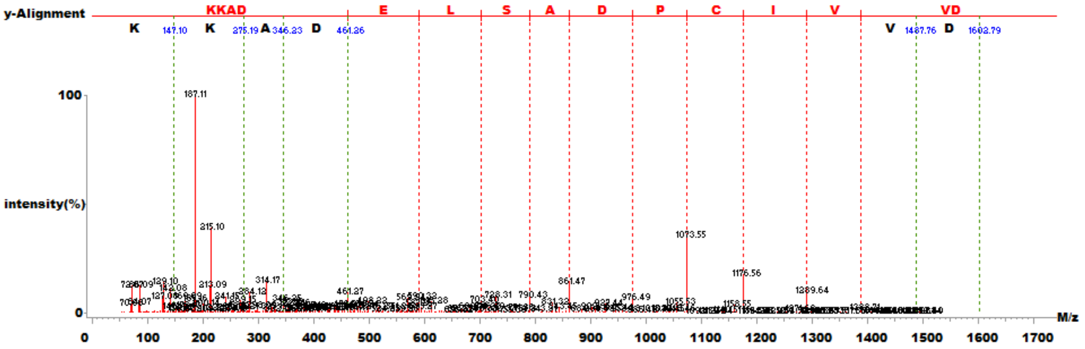


**Supplementary Figure 3.** (Continued)

$^{49}$DVVICPDASLEDAKK$^{63}$ + Cys-$SO_2H$ (C53)

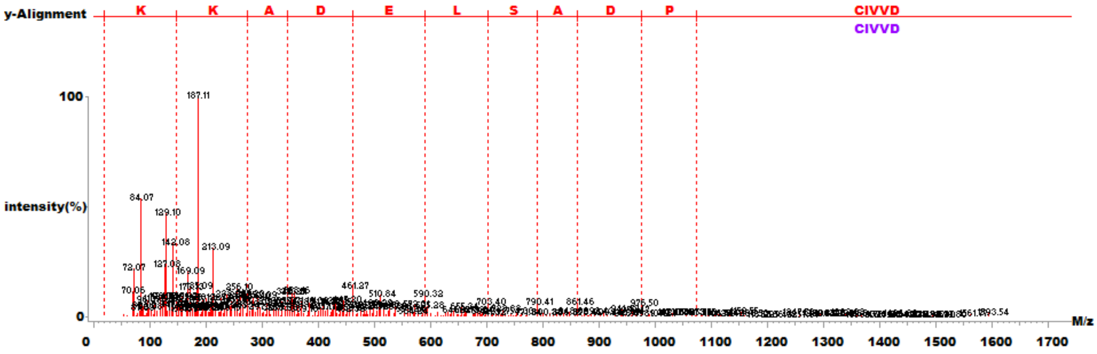


$^{49}$DVVICPDASLEDAKK$^{63}$ + Cys-$SO_3H$ (C53)

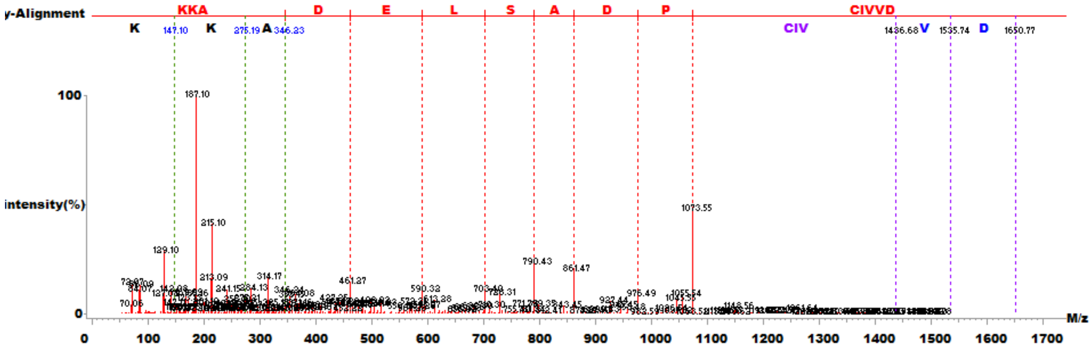


$^{49}$DVVICPDASLEDAKK$^{63}$ + Cys to Ser (C53)

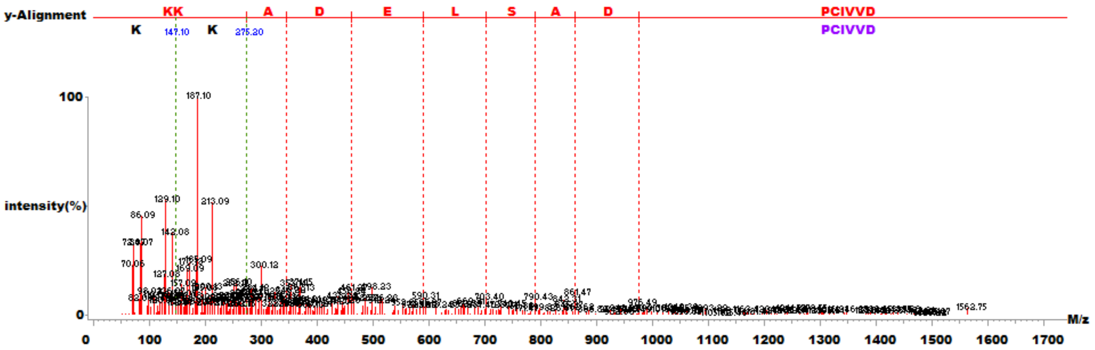


$^{49}$DVVICPDASLEDAKK$^{63}$ + Cys-$SO_2$-SH (C53)

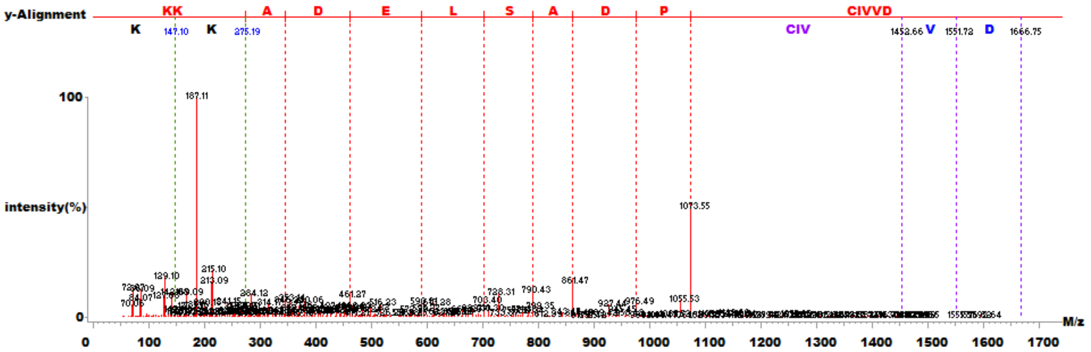


**Supplementary Figure 3.** (Continued)

a

| Native | 1-10 | 10-16 | 17-26 | 27-45 | 37-49 | 50-58 | 59-82 | 83-92 | 93-103 | 104-112 | 117-131 | 132-144 | 145-153 | 152-163 | 164-170 | 171-182 | 183-189 |
|---|---|---|---|---|---|---|---|---|---|---|---|---|---|---|---|---|---|
| 10 sec | 10 | 14 | 0 | 21 | 9 | 11 | 29 | 20 | 9 | 11 | 20 | 23 | 22 | 8 | 0 | 0 | 29 |
| 60 sec | 10 | 14 | 0 | 32 | 9 | 11 | 29 | 20 | 18 | 11 | 20 | 31 | 22 | 21 | 0 | 8 | 29 |
| 300 sec | 20 | 14 | 0 | 32 | 27 | 22 | 38 | 30 | 18 | 11 | 33 | 31 | 28 | 25 | 0 | 8 | 43 |
| 1800 sec | 20 | 14 | 0 | 32 | 27 | 22 | 38 | 30 | 27 | 17 | 33 | 38 | 28 | 25 | 0 | 8 | 50 |
| 10800 sec | 20 | 29 | 0 | 42 | 36 | 22 | 42 | 30 | 32 | 28 | 40 | 46 | 33 | 29 | 0 | 8 | 57 |

| oxidized | 1-10 | 10-16 | 17-26 | 27-45 | 37-49 | 50-58 | 59-82 | 83-92 | 93-103 | 104-112 | 117-131 | 132-144 | 145-153 | 152-163 | 164-170 | 171-182 | 183-189 |
|---|---|---|---|---|---|---|---|---|---|---|---|---|---|---|---|---|---|
| 10 sec | 10 | 14 | 0 | 21 | 0 | 11 | 25 | 20 | 9 | 11 | 20 | 31 | 22 | 17 | 0 | 8 | 43 |
| 60 sec | 10 | 14 | 0 | 26 | 0 | 11 | 29 | 20 | 18 | 11 | 20 | 31 | 22 | 17 | 0 | 8 | 43 |
| 300 sec | 10 | 14 | 0 | 32 | 8 | 22 | 38 | 20 | 18 | 11 | 27 | 31 | 28 | 25 | 0 | 8 | 57 |
| 1800 sec | 10 | 14 | 0 | 37 | 8 | 22 | 42 | 30 | 27 | 22 | 40 | 38 | 28 | 25 | 0 | 8 | 57 |
| 10800 sec | 20 | 29 | 0 | 53 | 8 | 22 | 46 | 50 | 45 | 44 | 47 | 46 | 33 | 42 | 0 | 8 | 57 |

0~10 (%)
11~20 (%)
21~30 (%)
31~40 (%)
41~50 (%)
51~ (%)

b

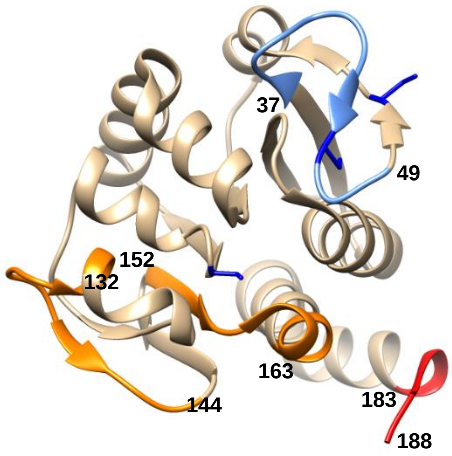


**Supplementary Figure 4. Identification of structural changes in DJ-1 under the oxidative stress employing HDX-MS.** Recombinant DJ-1 treated with and without $H_2O_2$ for 1 h, was incubated with $D_2O$ exchange buffer at 25ºC for various times upto 3 h and analyzed using nanoAcquity™ /ESI/MS. (a) Deuterium exchange rate (%) of native and oxidized DJ-1 was presented depending on $D_2O$ incubation time. Significant changed representative peptide were marked by colored arrows and no discernible changes by gray arrow. The corresponding deuterium exchange levels of each peptide in percent are given on the right. (b) Average deuterium exchange difference of oxidized DJ-1 compared to the native DJ-1 monomer structure (PBD ID code 4RKW).

a

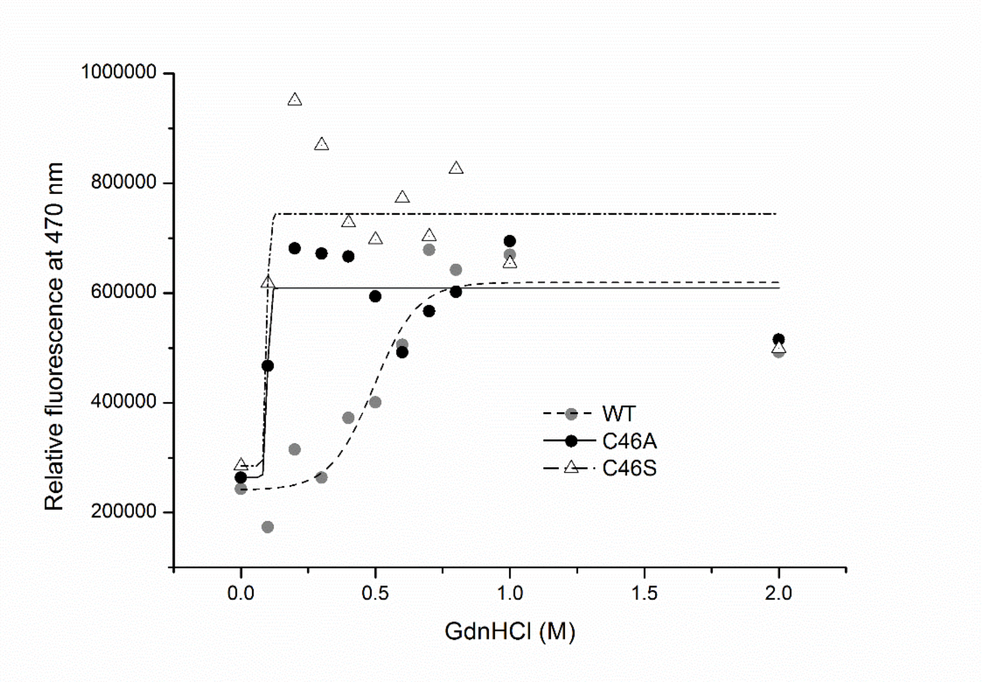


b

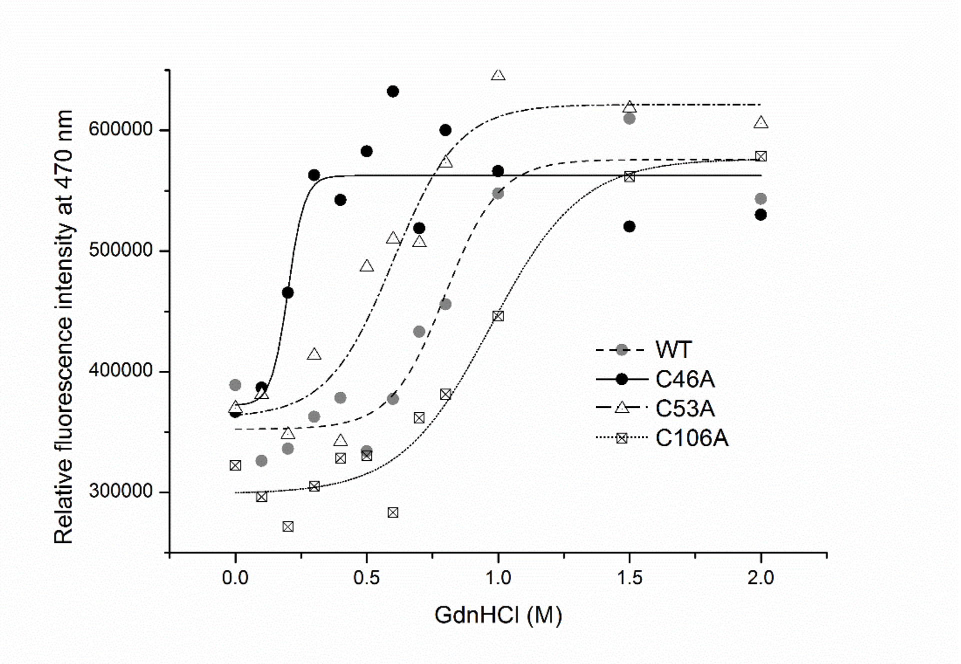


**Supplementary Figure 5. Protein stability and folding of DJ-1 proteins in response to denaturants.** Hydrophobic exposure during GdnHCl-induced unfolding was monitored by ANS fluorescence intensity at 470 nm. (a) WT, C46A and C46S mutant and (b) WT, C46A, C53A and C106A were incubated for 16 h with various concentrations of GdnHCl. Samples were prepared by mixing the protein with ANS stock solution to final molar ration of 75 : 1 (ANS : protein) and equilibrating in the dark for 30 min at R.T. Fluorescent emission intensity at 470 nm was measured using an excitation wavelength 380 nm. Data for each proteins were fitted with a Boltzmann curve using Origin 8.5.

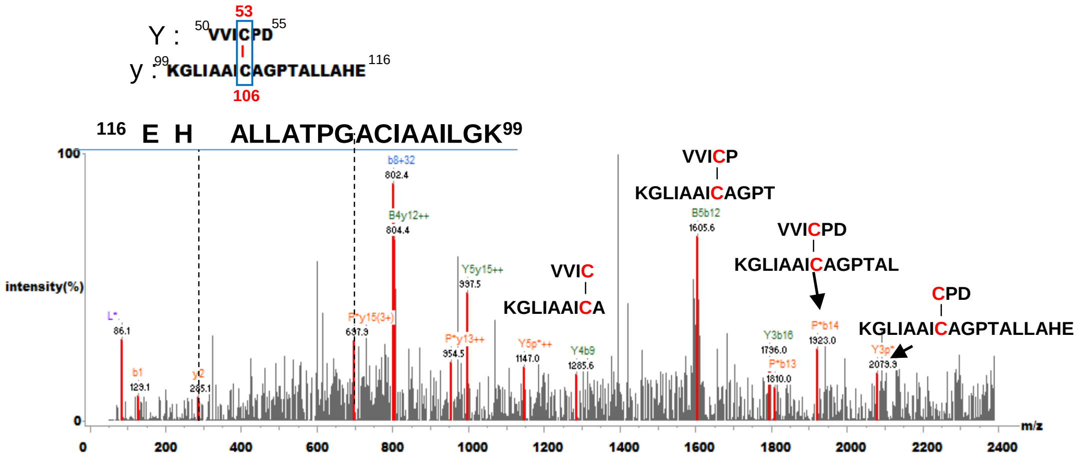


**Supplementary Figure 6.** Tandem mass spectrum of Cys53-Cys106 disulfide bond peptide in the $H_2O_2$-untreated C46A mutant band in Figure 6a. Silver-stained gels were cut out of the gel and analyzed by MS and DBond analysis was performed.

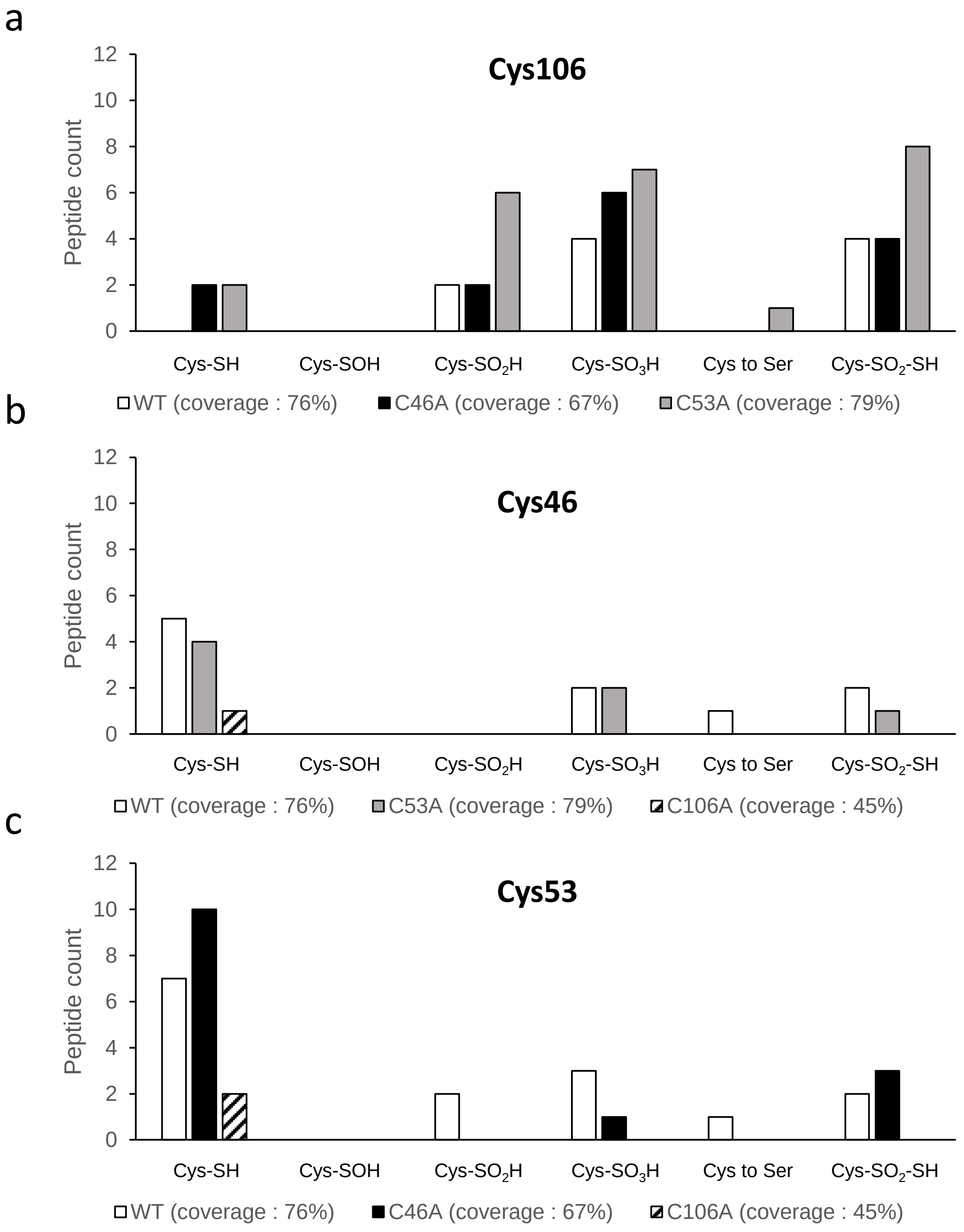


**Supplementary Figure 7.** Each cysteine influenced on the oxidation state of other cysteines with different modifications in Hela cells. (a-c) WT and Cys mutants of DJ-1 protein bands without $H_2O_2$ treatment in Figure 6a under reducing condition were examined by MS/MS analysis employing SEMSA strategy and Mascot search algorithm. The number of oxidized peptides of (a) Cys106, (b) Cys46 and (c) Cys53 in WT and Cys mutants of DJ-1 were presented with sample coverage. Various oxidative modifications such as sulfinic acid, sulfonic acid , conversion of Cys to Ser and  conversion of Cys to thiosulfonic acid ($\Delta m = +64$ Da) were found.

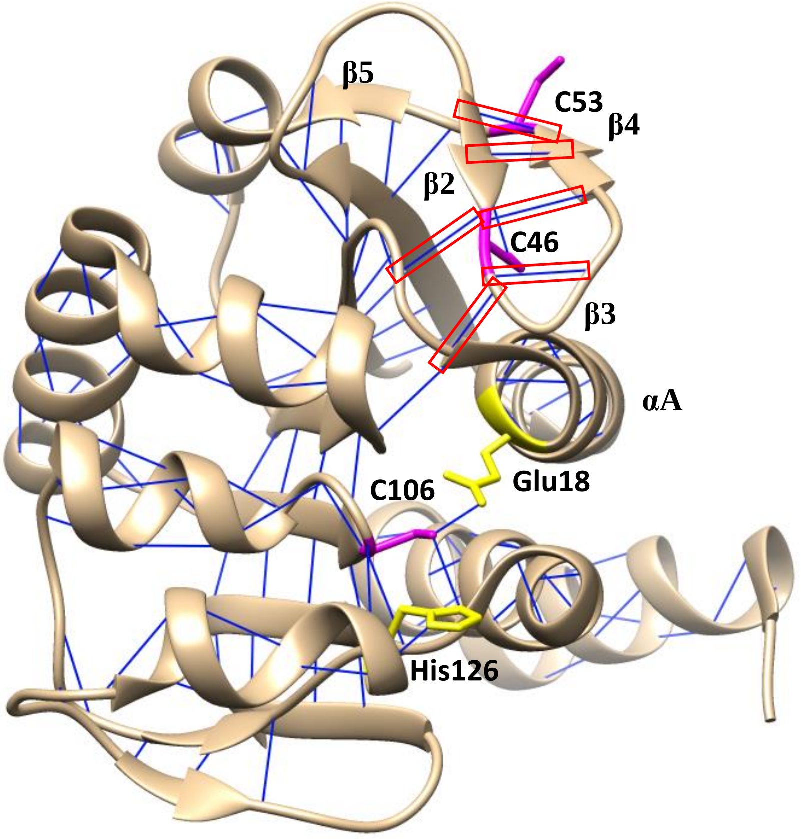


**Supplementary Figure 8.** Prediction of hydrogen-bonds in DJ-1 using UCSF Chimera program (PDB ID code : 4RKW). Cysteine residues are shown in magenta and Glu18, which regulate activity of Cys106, is shown in yellow. H-bonds existing in the long loops bearing β2-β4 sheet are marked with red square.
